# The Intrinsically Disordered Region of ExbD is Required for Signal Transduction

**DOI:** 10.1101/797811

**Authors:** Dale R. Kopp, Kathleen Postle

## Abstract

The TonB system actively transports vital nutrients across the unenergized outer membranes of the majority of Gram-negative bacteria. In this system, integral membrane proteins ExbB, ExbD, and TonB work together to transduce the protonmotive force (PMF) of the inner membrane to customized active transporters in the outer membrane by direct and cyclic binding of TonB to the transporters. A PMF-dependent TonB-ExbD interaction is prevented by 10-residue deletions within a periplasmic disordered domain of ExbD adjacent to the cytoplasmic membrane. Here we explored the function of the ExbD disordered domain in more detail. *In vivo* photo-cross-linking through sequential pBpa substitutions in the ExbD disordered domain captured five different ExbD complexes, some of which had been previously detected using in vivo formaldehyde crosslinking, a technique that lacks the residue-specific information that can be achieved through photo-cross-linking: 2 ExbB-ExbD heterodimers (one of which had not been detected previously), previously detected ExbD homodimers, previously detected PMF-dependent ExbD-TonB heterodimers, and for the first time, a predicted, ExbD-TonB PMF-*in*dependent interaction. The fact that multiple complexes were captured by the same pBpa substitution indicated the dynamic nature of ExbD interactions as the energy transduction cycle proceeded in vivo. In this study, we also discovered that a conserved motif, (V45, V47, L49, P50), within the disordered domain was required for signal transduction to TonB and to the C-terminal domain of ExbD and was the source of its essentiality.

**Importance:** The TonB system is a virulence factor for many Gram-negative pathogens including E-S-K-A-P-E pathogenic species *Klebsiella pneumoniae*, *Acinetobacter baumannii*, *and Pseudomonas aeruginosa*. Because the majority of protein-protein interactions in the TonB system occur in the periplasm, it is an appealing target for novel antibiotics. Understanding the molecular mechanism of the TonB system will provide valuable information for design of potential inhibitors targeting the system.

## INTRODUCTION

The TonB system is required for most Gram-negative bacteria to acquire vital nutrients such iron, vitamin B_12_, nickel, cobalt, copper, heme, maltodextrin, and sucrose across their outer-membrane barriers and is major virulence factor in many pathogenic Gram-negative bacteria (1, 2). Under iron-limiting conditions, such as the human serum, Gram-negative bacteria secrete iron-chelating compounds called siderophores that capture the iron with high affinity (3). However, because iron-siderophore complexes are too large, too scarce, and too important to passively diffuse through the outer-membrane porins, they require customized 22-stranded beta-barrel transporters called TonB-dependent transporters (TBDTs) to transition across the outer membrane (4, 5).

After the siderophore captures an iron atom extracellularly, the iron-siderophore complex binds to a TDBT with sub-nanomolar affinity, causing the ligand-loaded TBDT to signal on its periplasmic face that ligand is bound (6–9). Energy is required to release the iron-siderophore from the TBDT (10). Because the outer membrane and periplasmic space lack a sufficient energy source, Gram-negative bacteria transduce the energy from the cytoplasmic membrane proton motive force (PMF) to the TBDTs using the three known integral cytoplasmic membrane proteins of the TonB system--ExbB, ExbD, and TonB (11, 12).

In *Escherichia coli* K-12, there are six TBDTs that transport various sources of iron and one TBDT that transports vitamin B12. With a cellular ratio of 7 ExbB: 2 ExbD: 1 TonB, ExbB and ExbD harvest PMF and transmit it to TonB by a largely unknown mechanism (13). ExbB has three transmembrane domains (TMDs) with the majority of the protein residing in the cytoplasm [Fig. 1, (14, 15)]. In contrast, ExbD and TonB are anchored by single N-terminal TMDs with the bulk of their residues occupying the periplasm (16, 17). ExbB, ExbD, and TonB also share homology through their TMDs with proteins TolQ, TolR, and TolA of the Tol system (respectively), which is important for outer membrane integrity and cell division (18, 19). Both systems are energized by PMF, suggesting that they have a common means of PMF utilization that plays out as different roles in cell physiology due to different protein contacts made by their soluble domains (20, 21).

**Figure 1:**
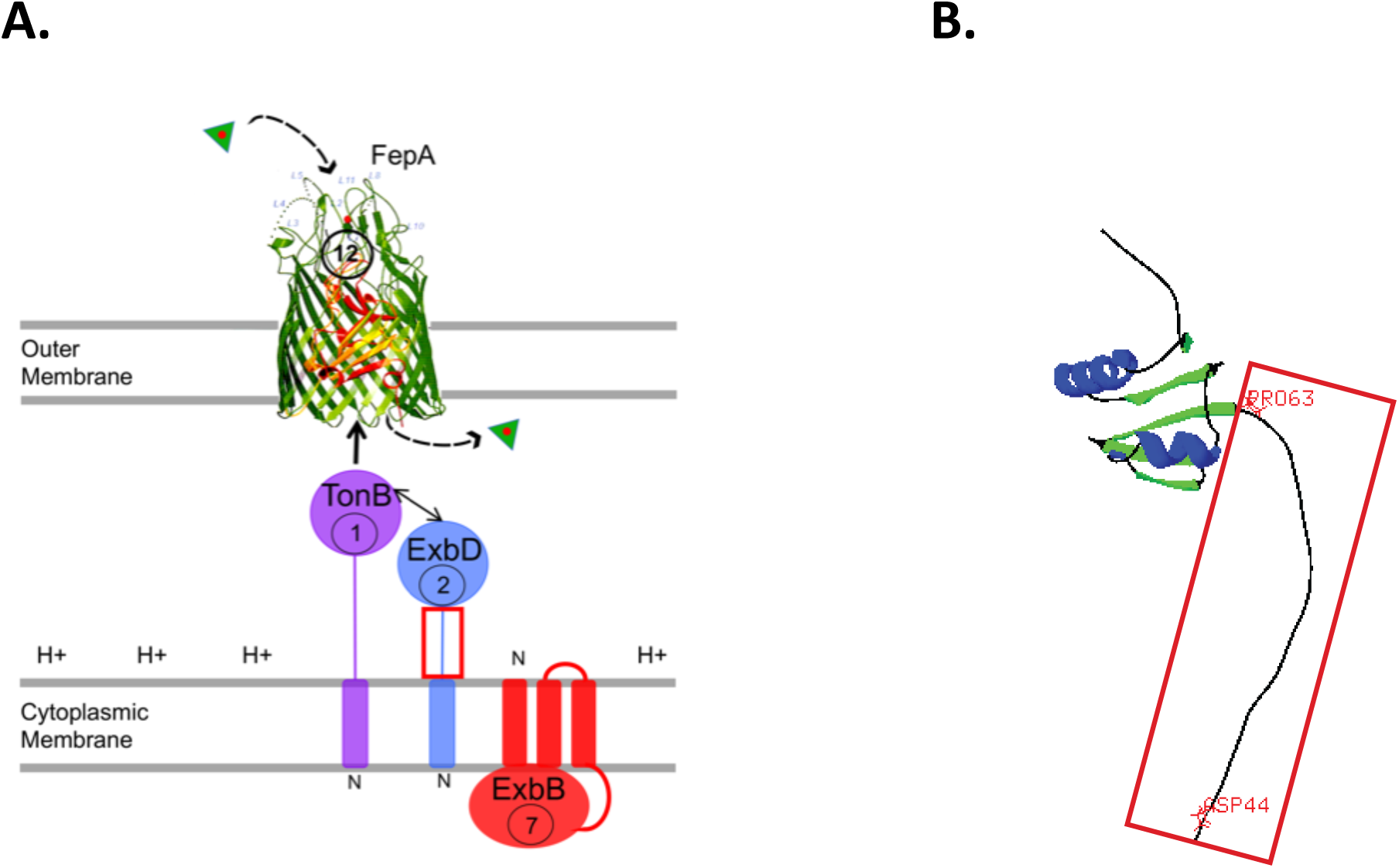
(A) TonB-dependent energy transduction. The iron-bearing siderophore enterochelin (triangles) binds to the TBDT FepA with sub-nanomolar affinity (71). This high affinity necessitates a mechanism to release the siderophore into the periplasmic space. In the presence of cytoplasmic membrane proteins TonB, ExbB, ExbD, and the cytoplasmic membrane proton motive force (PMF), the Fe-enterochelin is actively transported across the outer membrane through FepA. Numbers indicate the ratio of the proteins in *E. coli* under iron-limiting conditions (13). Because TonB is limiting, there must be a cycle during which TonB binds and releases TBDTs. The Stages in the cycle have been studied, resulting in a model where ExbB acts as a scaffold to manage an essential PMF-dependent interaction between TonB and ExbD that subsequently positions TonB for correct interaction with FepA. The red box around ExbD identifies the ExbD disordered region which is the topic of this study. Subsequent transport of enterochelin across the cytoplasmic membrane does not require the TonB system [adapted from (24)]. (B) The NMR solution structure of ExbD periplasmic domain residues 44-141 [NCBI PDB ID: 2PFU; (37)]. The alpha helices and beta strands are colored in blue and green, respectively. The ExbD disordered region is boxed in red. The first residue following the ExbD TMD, D44, and last residue, P63 of the ExbD disordered region are labeled with the side chain displayed. This image was generated using Swiss-PdbViewer [https://spdbv.vital-it.ch/(37)].

The TBDTs greatly outnumber the molecules of TonB in a cell, leading to competition between TBDTs for “energized” TonB (13, 22). TonB must thus undergo a cyclic pattern of binding and release from TBDT to TBDT, referred to as the TonB energization cycle (6, 23, 24). TonB binds to TBDTs both *in vitro* and *in vivo* whether or not it is “energized”, and therefore, it must be correctly configured during an energy transduction cycle to enable productive interaction with the TBDT (24–28). Both ExbB and ExbD are required to correctly configure TonB to energize active transport across the outer membrane--ExbB indirectly, and ExbD directly (15, 29, 30).

ExbB serves as a scaffolding protein for ExbD and TonB assembly, without which they are both proteolytically unstable, and mediates ExbD-TonB interactions (15, 31–33). ExbD-TonB interactions, or lack thereof, define our model for the initial Stages of the TonB energization cycle (24, 34). In Stage I, ExbD and TonB homodimers exist in separate complexes with ExbB homotetramers, but do not interact. To transition to Stage II, ExbB tetramers bring ExbD and TonB homodimers together in a single complex without a requirement for PMF. The PMF-*in*dependent ExbD-TonB interaction has only been detected indirectly by ExbD-mediated protection of the N-terminal ∼155 residues of TonB from exogenously added proteinase K in spheroplasts that lack PMF (6, 15, 35). In the subsequent presence of PMF, Stage III, the ExbD and TonB homodimers swap partners to form ExbD-TonB heterodimers. Only in this Stage can the PMF-dependent ExbD-TonB interaction can be captured by formaldehyde cross-linking *in vivo* (29). The only TMD among the cytoplasmic membrane proteins of ExbB, ExbD, and TonB that can participate in response to PMF is the ExbD TMD, with its D25 residue (15, 29, 36).

ExbD has five structurally distinct domains: a cytoplasmically localized N-terminus (residues 1-22), a predicted TMD (residues 23-43), a disordered periplasmic domain [44-63], a structured periplasmic domain (residues 64-133) and a flexible C-terminus (residues 134-141) (37). Experiments using *in vivo* disulfide cross-linking of engineered Cys residues in the ExbD and TonB distal periplasmic domains are consistent with the three-Stage model. The same ExbD residues in the region of 92-121 that form a homodimeric interface are almost the exact same residues that form ExbD-TonB heterodimers through the TonB periplasmic domain (38). Also, in a PMF-unresponsive ExbD mutant, ExbD (D25N), ExbD-TonB disulfide cross-links are either absent or significantly reduced (39).

A 10 residue-deletion scanning mutagenesis of ExbD reveals that deletions within ExbD periplasmic residues 62-141 stall TonB energization in Stage I, whereas deletions largely encompassing the disordered domain of ExbD [42-61] stall TonB energization in Stage II, unable to proceed to Stage III (35). However, the mechanistic details of how the ExbD disordered domain participates in energy transduction have remained unclear.

Here we discovered that a novel motif, ΨXΨXLP (Ψ = hydrophobic-branched residues; X = non-hydrophobic residues) encompassing ExbD residues V45, V47, L49 and P50 and located adjacent to the ExbD TMD, is essential for ExbD signal transduction to its periplasmic C-terminus and to TonB. This motif within the disordered domain of ExbD is required for the transition from Stage II to Stage III and could account for crosstalk between the TonB and Tol systems because it is conserved in TolR. As might be predicted from its lack of structure, the disordered domain is highly dynamic, making contacts with TonB, another ExbD and two different contacts with ExbB. With respect to TonB contacts, the disordered domain forms the PMF-dependent complex previously visualized by formaldehyde cross-linking. Importantly, it also forms the long-predicted PMF-*in*dependent complex between ExbD and TonB, visualized for the first time without requiring collapse of PMF, thus indicating it is part of the normal energy transduction cycle. This study defines the role of the ExbD disordered domain and makes clear the dynamic character of protein-protein interactions that occur during TonB-dependent energy transduction.

## RESULTS

### Residues 44-53 in the ExbD disordered domain are essential for TonB system activity

In our previous study, deletions that encompassed most of the disordered domain of ExbD protein, (Δ42-51) and (Δ52-61), were completely inactive because they eliminated the ability of its partner protein, TonB, to undergo essential conformational changes in response to change in PMF (35). The effects of these deletions could have been due to structural distortions. To circumvent that issue, here we engineered and characterized ExbD mutagenized with blocks of Alanine substitutions to identify functionally important residues within the disordered domain (Fig. 1B): ExbD (44-48Ala), ExbD (49-53Ala), ExbD (54-58Ala), and ExbD (59-63Ala). TonB system activity was determined by measuring the initial transport rates of two different iron-siderophores, ^55^Fe-enterochelin and ^55^Fe-ferrichrome to identify any TonB-dependent transporter-specific activities (7, 40).

Both ExbD (54-58Ala) and ExbD (59-63Ala) transported ^55^Fe-enterochelin or ^55^Fe-ferrichrome at rates equivalent to the wild-type ExbD encoded on pKP999 (pExbD), identifying them as being in the mutation-tolerant half of the disordered domain. In the mutation-intolerant half, neither ExbD (44-48Ala) nor ExbD (49-53Ala) were able to transport either siderophore. ExbD (59-63Ala) required significantly more sodium propionate (20 mM) to achieve chromosomal ExbD levels, suggesting that it was important for proteolytic stability (Fig. 2). These results excluded residues 54-63 as functionally important for the TonB system and eliminated the possibility of siderophore-specific effects.

**Figure 2:**
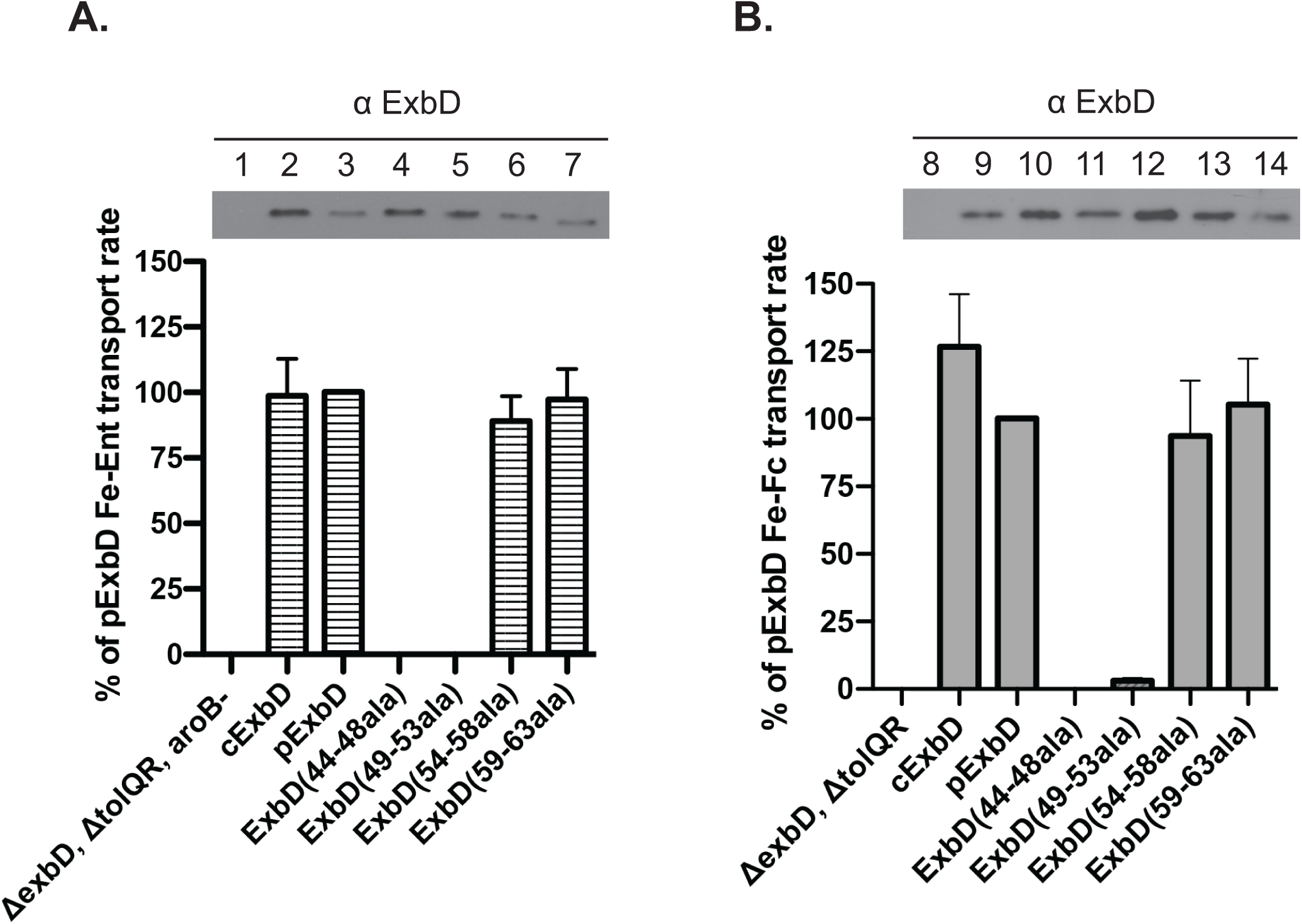
Block Alanine substitutions of ExbD residues 44-48 and 49-53 prevent TonB system activity. (A) Initial rate of ^55^Fe-enterochelin transport and corresponding steady-state expression levels of the ExbD proteins. Strains were grown to mid-exponential phase and assayed as described in Methods and Materials. The percentages of activity were calculated relative to the initial rate of plasmid-encoded ExbD, pExbD (pKP999), expressed at chromosomal levels in KP1526 (Δ*exbD*, Δ*tolQR, aroB*). Assays were performed in triplicate and carried out at least twice for each strain. The slopes were determined by a linear regression fit of scintillation counts per minute. The error bars indicate mean ± SEM for a representative triplicate set. The steady-state ExbD monomer levels at the time of assay were determined by immunoblot analysis (top). To avoid competition from enterochelin, the only siderophore synthesized by *Escherichia coli* K12 and transported by FepA (Fig. 1), *aroB* strains were used. cExbD designates chromosomally expressed ExbD from the wild-type strain, KP1270 (W3110, *aroB*). ExbD Alanine substitution mutants are expressed from plasmids in KP1526 (Δ*exbD*, Δ*tolQR, aroB*). (B) Initial rate of ^55^Fe-ferrichrome transport and corresponding steady-state expression levels of ExbD proteins. The assay was performed and analyzed identically as in (A) using the siderophore, ferrichrome, which is transported through the TonB-dependent transporter, FhuA. The percentages of activity were calculated relative to the initial rate of plasmid-encoded ExbD, pExbD (pKP999) expressed at chromosomal levels in RA1045 (Δ*exbD*, Δ*tolQR*). cExbD designates chromosomally expressed ExbD from the wild-type strain, W3110. All ExbD Alanine substitution mutants are expressed from plasmids in RA1045 (Δ*exbD*, Δ*tolQR*). The plasmid identities and concentrations of propionate for chromosomal expression are listed in Table S1.

Previously, ExbD (Δ42-51) and ExbD (Δ52-61) were unable to form the formaldehyde-cross-linked ExbD-TonB complex that typifies Stage III (35). Consistent with that, the inactive ExbD (44-48Ala) and ExbD (49-53Ala) had the same cross-linking profile in the presence and absence of TonB, indicating that they could not form that Stage III complex. In contrast, for the active ExbD (54-58Ala), and ExbD (59-63Ala), the absence of TonB led to the absence of the formaldehyde cross-linked ExbD-TonB complex (Fig. S1).

To test whether the absence of the PMF-dependent ExbD-TonB complex was due to the ability of ExbD (44-48Ala) and ExbD (49-53Ala) to somehow induce proton leakage, we measured the rate of PMF-dependent ethidium bromide efflux through the AcrA/AcrB/TolC complex (41, 42). None of the block Ala mutants came close to PMF leakage caused by 15 µM CCCP, which nonetheless still supports 100% ^55^Fe-ferrichrome transport (data not shown) (Fig. S2A). Because PMF was adequate for ^55^Fe-ferrichrome transport, the reason for the inactivity of the block Ala substitutions must lie elsewhere.

Expression of ExbD (44-48Ala) did, however, cause a small but statistically significant increase in ethidium bromide accumulation compared to pExbD (Fig. S2B). The means by which ExbD (44-48Ala) caused this low level of proton leakage was unclear, especially since the Δ*exbD* strain with *tolQR* present did not detectably leak protons (data not shown). Because there is crosstalk between the TonB system and the Tol-Pal system (43), and because RA1045 (Δ*exbD*, Δ*tolQR*) showed the same degree of proton leakage as ExbD (44-48Ala), we speculate that proton leakage could have been induced by TolA in the absence of TolQR or a functional ExbBD.

### *In vivo* photo-cross-linking identifies characteristic PMF-dependent and novel PMF-*in*dependent ExbD-TonB complexes

Disordered domains of proteins are often regions of interaction with other proteins (44). ExbD and TonB, both of which contain regions of disorder, form a formaldehyde cross-link, but the residues involved are unknown (29). To identify contacts made by the ExbD disordered domain, individual amber mutations were constructed at codons 44-63. The photoreactive, non-natural amino acid, *p*-benzoyl-L-phenylalanine (*p*Bpa) was incorporated at the site of each amber mutation and used to covalently capture protein interactions within 3 angstroms upon exposure to ultraviolet (UV) light (45, 46). The advantage of this approach over formaldehyde cross-linking is that interactions through defined sites can be investigated, while leaving the positions of residues in binding partners unconstrained.

To characterize ExbD-TonB interactions made through the ExbD disordered domain, we chose ExbD (S54pBpa) and ExbD (R59pBpa) as representatives for photo-cross-linking. Both variants were active in iron transport assays and shared representative complexes with the rest of the pBpa substitution mutants (shown below). On immunoblots developed with anti-TonB monoclonal antibodies, two different ExbD-TonB complexes were captured by both variants, a doublet at ∼55 kDa and a single band at ∼46 kDa (Fig. 2, lanes 4 and 8). Because a C-terminally added His_6_ epitope tag on ExbD caused each of the complexes to migrate at a higher apparent mass equivalent to that of the added His_6_ tag, all the bands consisted of TonB complexed with full-length ExbD (Fig. 2, lanes 11 and 12). No complexes were detected in a strain where the *tonB* gene was deleted (Fig. 2, lane 7).

The sole potentially proton-responsive amino acid in the TonB system, Asp 25, resides in the TMD of ExbD [Fig.1; (15, 36)]. ExbD (D25N) is a well-characterized inactive mutant which causes the TonB energization cycle to stall at Stage II and is therefore useful for differentiating between PMF-dependent and -*in*dependent ExbD interactions (29, 35, 47). Complexes decreased or eliminated by ExbD (S54pBpa) or ExbD (R59pBpa) in concert with the D25N mutation identified the ∼55 kDa doublet as being PMF-dependent (Fig. 2, lanes 5 and 9). Consistent with that observation, the same complexes were also eliminated by treatment with the protonophore cyanide *m*-chlorophenylhydrazone (CCCP) (Fig. 2, lanes 6 and 10). Although the reason for the doublet was not clear, it appeared to represent two different PMF-dependent sites of contact made by ExbD (S54pBpa) and ExbD (R59pBpa) with TonB.

In contrast, the abundance of ∼46 kDa ExbD-TonB complex was undiminished by the presence of the ExbD D25N mutation or by CCCP treatment, indicating that it formed independently of whether or not PMF was present. The ExbD D25N variant generated somewhat higher levels of the ∼46 kDa complex, consistent with the idea that it stalls TonB at Stage II, where such complexes build up (34). These data demonstrated that loss of PMF caused a significant change in the sites through which ExbD disordered domain residues contact TonB—enough difference that it resulted in a change in the apparent mass of complexes from ∼55 kDa to ∼46 kDa.

We knew that ExbD and TonB made *in vivo* contacts through their extreme C-termini (38, 39). We did not know that additional ExbD contacts occurred so close to the ExbD (and, presumably, TonB) TMDs (Fig. 1A,B).

### Most of the pBpa substitutions in the ExbD disordered domain capture some form of TonB contact

To understand the full extent of interactions between the ExbD disordered domain and TonB, *in vivo* photo-cross-linking was carried out through pBpa substitutions at ExbD codons 44-63. Most pBpa substitutions in the region from 44-52 supported significantly reduced TonB-dependent ^55^Fe transport activity (< 25% wild-type levels), consistent with the results from the block Ala substitutions which essentially eliminated activity (compare Figs.2 and 4A). Substitutions at residues 53-63 all supported at least 50% activity except for ExbD (R59pBpa) which also supported only 25% activity. Several single pBpa substitution mutants from residues 54-63 were less active than ExbD (54-58Ala) and ExbD (59-63Ala) (compare Fig. 2A with Fig. 4A, right panel). The bulky pBpa side chain was likely responsible for the discrepancy, possibly alluding to the importance of conformational flexibility or lock and key fit for these residues. Nonetheless, except for ExbD (V45pBpa) and ExbD (V47pBpa), all the pBpa substitutions within the ExbD disordered domain had detectable levels of activity. Therefore, the complexes captured by these substitutions at least identified TonB proteins within ∼3angstroms of the pBpa moiety (46), and likely reflected functional interactions that occurred during a TonB energization cycle.

**Figure 4:**
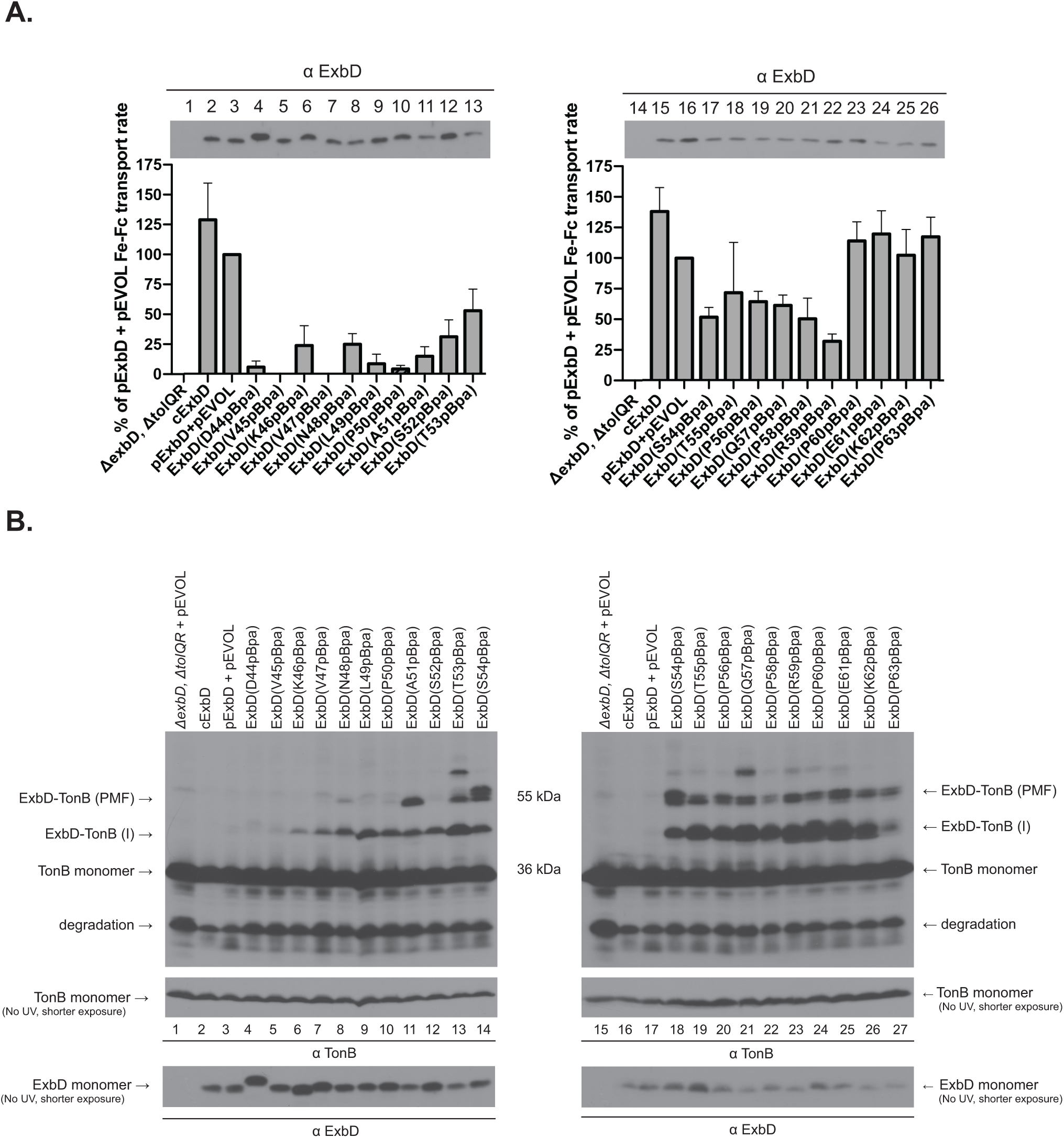
Virtually all the ExbD (pBpa) substitutions in the ExbD disordered region capture PMF-dependent or PMF-*in*dependent TonB complexes. (A) Initial rate of ^55^Fe-ferrichrome transport for ExbD (pBpa) substitutions at residues 44-54 (left panel) and residues 54-63 (right panel) with the corresponding steady-state expression levels of ExbD proteins at the time of assay above each panel. Plasmid-encoded ExbD pBpa substitutions were co-expressed with plasmid pEVOL in RA1045 (Δ*exbD*, Δ*tolQR*). Strains were grown to mid-exponential phase and assayed as described in Methods and Materials. The percentages of activity were calculated relative to the initial rate for ^55^Fe-ferrichrome transport of plasmid-encoded ExbD, pExbD (pKP999). Assays were performed in triplicate and carried out at least twice for each strain. The slopes were determined by a linear regression fit of scintillation counts per minute. The error bars indicate mean ± SEM for a representative triplicate set. cExbD designates chromosomally expressed ExbD from the wild-type strain, W3110. The plasmid identities and concentrations of propionate for chromosomal expression are listed in Table S1. (B) *In vivo* photo-cross-linking of pBpa substitutions for ExbD residues 44-54 (left panel) and residues 54-63 (right panel) on an anti-TonB immunoblot. Note that ExbD S54 (pBpa) is included on both immunoblots in lanes 14 and 18 for comparison of levels of expression. On the left and right, positions of a PMF-dependent ExbD-TonB complex [ExbD-TonB (PMF)], a PMF-*in*dependent complex [ExbD-TonB (I)], monomeric TonB, and a TonB degradation product are shown. Mass markers are indicated in the center. The steady-state levels of ExbD and TonB monomers (bottom) from the same experiment were determined by immunoblot analysis of control samples that were not exposed to UV light.

The PMF-dependent ExbD-TonB complexes formed starting at approximately ExbD residue A51(pBpa) and continuing through P63 (pBpa), with all the pBpa substitutions except S52(pBpa) participating (Fig. 4B). Consistent with the intrinsically disordered nature of this region and the already known changes in protein-protein interactions that must occur during an energy transduction cycle, there was not a strong pattern of interaction suggestive of an *α*-helix or *β*-sheet (24, 35, 37).

We had previously observed a PMF-dependent formaldehyde-cross-linked ExbD-TonB complex, but did not know through which residues of the ExbD periplasmic domain it occurred (29). The ExbD disordered domain contains only three formaldehyde-cross-linkable residues: K46, R59, and K62 (48). ExbD (K46pBpa) did not capture the PMF-dependent ExbD-TonB interaction via photo-cross-linking, indicating that the PMF-dependent formaldehyde cross-linked complex did not form through that residue (Fig. 4B, left panel). Similarly, because ExbD (59-63Ala), which eliminated formaldehyde-crosslinkable residues R59 and K62, still captured the PMF-dependent formaldehyde-cross-linked ExbD-TonB complex (Fig. S1), the idea that the ExbD disordered domain participated in the previously observed PMF-dependent formaldehyde cross-link to TonB could be ruled out.

The PMF-*in*dependent ExbD-TonB interaction characterized in Fig. 3, was captured by the majority of the residues within the ExbD disordered domain (Fig. 4B). Because the abundance of complex was fainter at either end of the range of pBpa substitutions, it could be that the ExbD disordered domain constituted the PMF-*in*dependent interface. Interestingly, the ExbD disordered residues that trapped the PMF-*in*dependent interactions were similar to an ExbD interface predicted to interact with TonB by phage-panning experiments (49).

**Figure 3:**
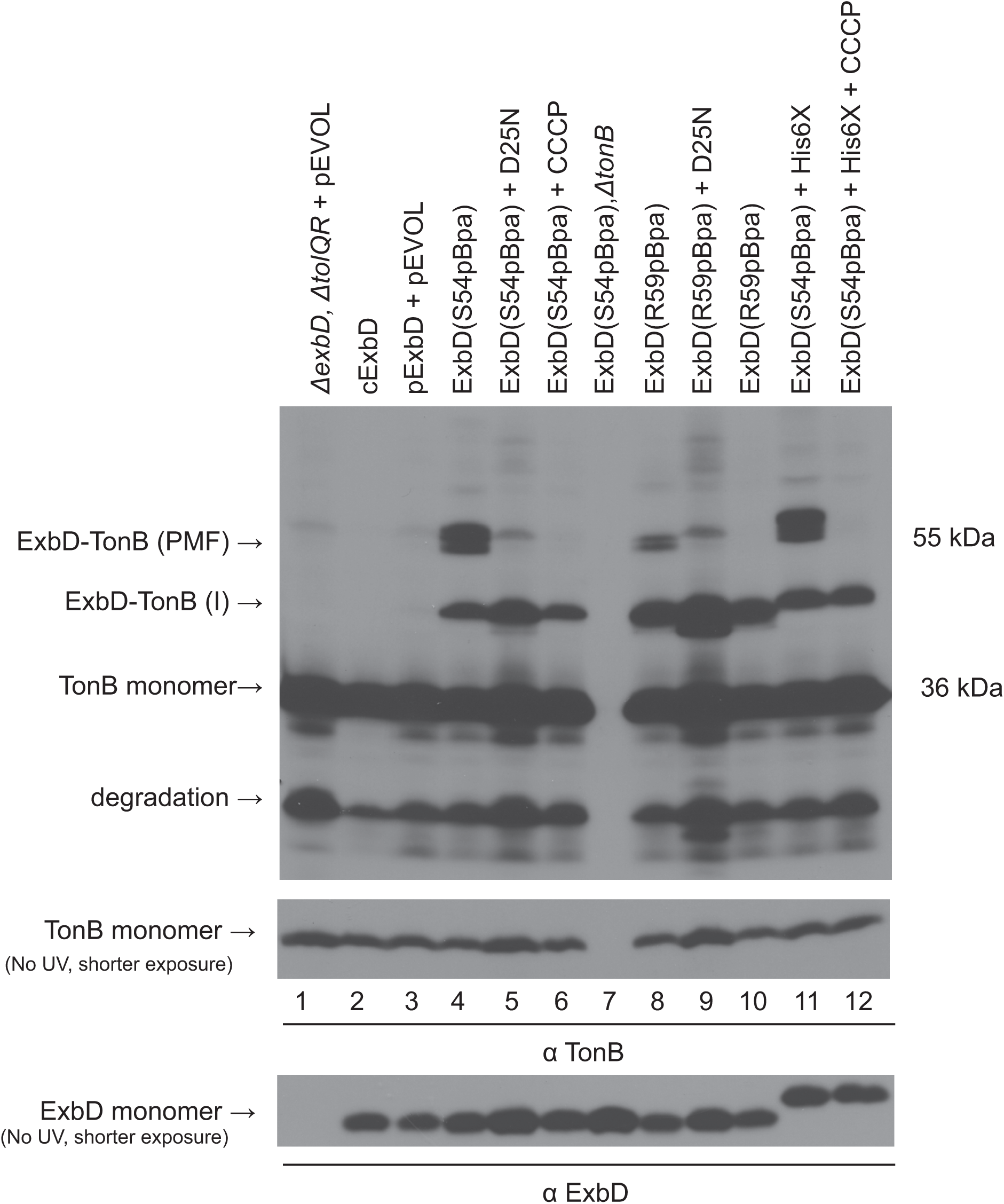
Both ExbD-TonB PMF-dependent and PMF-independent complexes can be identified by *in vivo* photo-cross-linking. Plasmid-encoded ExbD (pBpa) substitutions were co-expressed with pEVOL in strain RA1045 (Δ*exbD*, Δ*tolQR*). At mid-exponential phase, strains were subjected to UV irradiation for photo-cross-linking as described in Materials and Methods. Equivalent numbers of cells were precipitated with TCA and the ExbD-TonB complexes were visualized on immunoblots of 13% SDS-polyacrylamide gels by anti-TonB monoclonal antibodies. cExbD designates chromosomally expressed ExbD from the wild-type strain, W3110. The parent plasmid strain pExbD (pKP999) and all ExbD (pBpa) substitutions were expressed in RA1045 (Δ*exbD*, Δ*tolQR*), except for ExbD (S54pBpa) Δ*tonB*, which was expressed in KP1509 (Δ*exbD*, Δ*tolQR, ΔtonB*). D25N is a mutation in the ExbD TMD. The protonophore CCCP was added prior to UV exposure. His6X refers to ExbD (S54pBpa) with a C-terminal His_6_ tag. On the left, positions of a PMF-dependent ExbD-TonB complex [ExbD-TonB (PMF], a PMF-*in*dependent complex [ExbD-TonB (I)], monomeric TonB, and a TonB degradation product are shown. Mass markers are indicated on the right. Steady state ExbD and TonB monomer levels from the same experiment were determined by immunoblot analysis of control samples that were not exposed to UV light (bottom). The plasmid identities and concentrations of propionate for chromosomal expression are listed in Table S1.

In addition to the ExbD-TonB PMF-dependent complex and ExbD-TonB PMF-*in*dependent complex, there was also an unknown ∼62 kDa complex trapped via photo-cross-linking by ExbD (T53pBpa) and ExbD (Q57pBpa) (Fig. 4B). We do not know the identity of the complex except that it certainly contains both ExbD and TonB.

### The ExbD disordered domain interacts with another ExbD and with ExbB

ExbD also formaldehyde cross-links *in vivo* into a single homodimeric complex (∼ 30 kDa) and a single heterodimeric complex with ExbB (∼41 kDa) through unknown residues (29, 34). To determine if the ExbD disordered domain formed similar complexes, we used ExbD (D44pBpa) and (S54pBpa) as examples (Fig. 5A, B). In Fig. 5A, ExbD (D44 pBpa) formed complexes with apparent masses of ∼39 kDa and ∼43 kDa, making them both, based on their apparent masses, candidates to be potential ExbD-ExbB complexes. Surprisingly, the addition of a C-terminal His_6_ epitope tag to ExbB caused both complexes to shift to a higher mass consistent with the added His_6_ tag, indicating that both were ExbD-ExbB complexes. The presence of two different ExbD-ExbB complexes indicated that either or both proteins were undergoing conformational changes that influenced the nature of the contacts between them during the energy transduction cycle.

**Figure 5:**
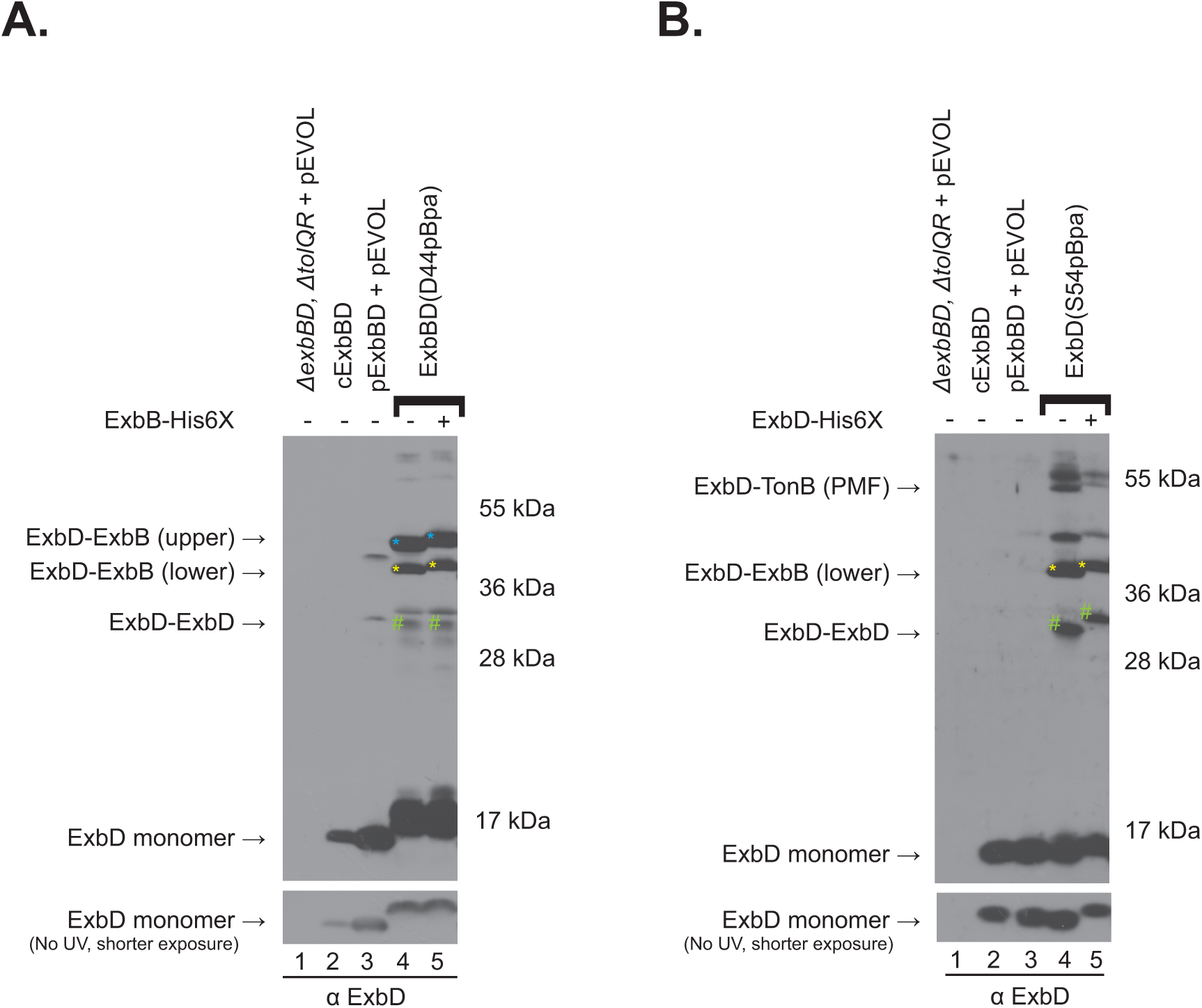
Two different ExbD-ExbB complexes and an ExbD homodimer are identified by *in vivo* photo-cross-linking. (A) *In vivo* photo-cross-linking of a representative ExbD photo-cross-linkable mutant, ExbD (D44pBpa), in the presence of ExbB with (+) and without (-) a C-terminal His_6_ epitope tag (His6X). At mid-exponential phase, strains were subjected to UV irradiation for photo-cross-linking as described in Materials and Methods. Equivalent numbers of cells were precipitated with TCA and the ExbD complexes were visualized on immunoblots of 13% SDS-polyacrylamide gels with anti-ExbD polyclonal antibodies. cExbD designates chromosomally expressed ExbB and ExbD from the wild-type strain, W3110. Plasmid pExbBD (pKP1657) and its corresponding ExbD (D44pBpa) variants were expressed with pEVOL in RA1017 (W3110, *ΔexbBD::kan, ΔtolQRA*). ExbD is overexpressed from plasmid pExbBD and its corresponding ExbD (D44pBpa) variant in the absence of inducer relative to chromosomally ExbD expression under the conditions of this assay. Positions of ExbD-ExbB (upper)—blue asterisk on immunoblot, ExbD-ExbB (lower)—yellow asterisk on immunoblot, likely ExbD-ExbD homodimers–green pound symbol on immunoblot, and ExbD monomer are indicated on the left. pExbBD expressed with pEVOL (lane 3) forms two unidentified complexes that were not ExbD residue-specific (unlabeled). Mass markers are indicated on the right. Steady-state ExbD monomer levels from the same experiment were determined by immunoblot analysis of control samples that were not exposed to UV light. The plasmid identities and concentrations of propionate for chromosomal expression are listed in Table S1. (B) *In vivo* photo-cross-linking of ExbD (S54pBpa), with (+) and without (-) a C-terminal His_6_ tag (His6X) using the same procedure as in (A). Plasmid pExbD (pKP999) and a corresponding variant, ExbD (S54pBpa) were expressed with pEVOL in RA1045 (Δ*exbD*, Δ*tolQR*). On the left, positions of a PMF-dependent ExbD-TonB complex [ExbD-TonB (PMF], ExbD-ExbB, an ExbD homodimer, and monomeric ExbD are shown. Mass markers are indicated on the right. Steady-state ExbD monomer levels from the same experiment were determined by immunoblot analysis of control samples that were not exposed to UV light (bottom). The plasmid identities and concentrations of propionate for chromosomal expression are listed in Table S1.

In Fig. 5B, ExbD (S54pBpa) formed an ∼31 kDa complex, consistent with an ExbD homodimer. Addition of a C-terminal His_6_ tag caused a decrease in the migration of the ∼31 kDa complex consistent with the presence of two His_6_ epitope tags, confirming that identification. (Fig. 5B). ExbD (S54pBpa) also formed the ∼ 46 kDa ExbD-ExbB complex and, as expected, the ExbD-TonB PMF-dependent and -*in*dependent complexes.

### The majority of pBpa substitutions in the ExbD disordered domain capture contacts with ExbB as well as ExbD

To understand the full extent of interaction between the ExbD disordered domain and itself or ExbB, we analyzed in vivo photo-cross-linking by the series of ExbD (pBpa) substitutions by immunoblot with anti-ExbD antibody. As expected, the ExbD-TonB PMF-dependent and -*in*dependent complexes could be discerned, but they were less confidently identified due to the presence of other complexes in the same region of the immunoblot, indicating that ExbD interacted with additional proteins, as seen in Fig. 5.

The ExbB-ExbD (upper, ∼43 kDa) complex was prominently formed by residues from the beginning of the ExbD disordered domain D44pBpa through P50pBpa, contacts that placed the region adjacent to the predicted ExbD TMD, close to the small periplasmic regions of ExbB, which mainly occupies the cytoplasm (Fig.1). Based on results with anti-TonB antibodies (Fig. 4 lanes 5-10), it is likely that ExbD pBpa 46-50 lanes (Fig. 6, lanes 5-10) also contain ExbD-TonB PMF-*in*dependent complex, which runs similarly to the ExbB-ExbD (upper, ∼43 kDa) complex and is therefore difficult to separately identify.-

**Figure 6:**
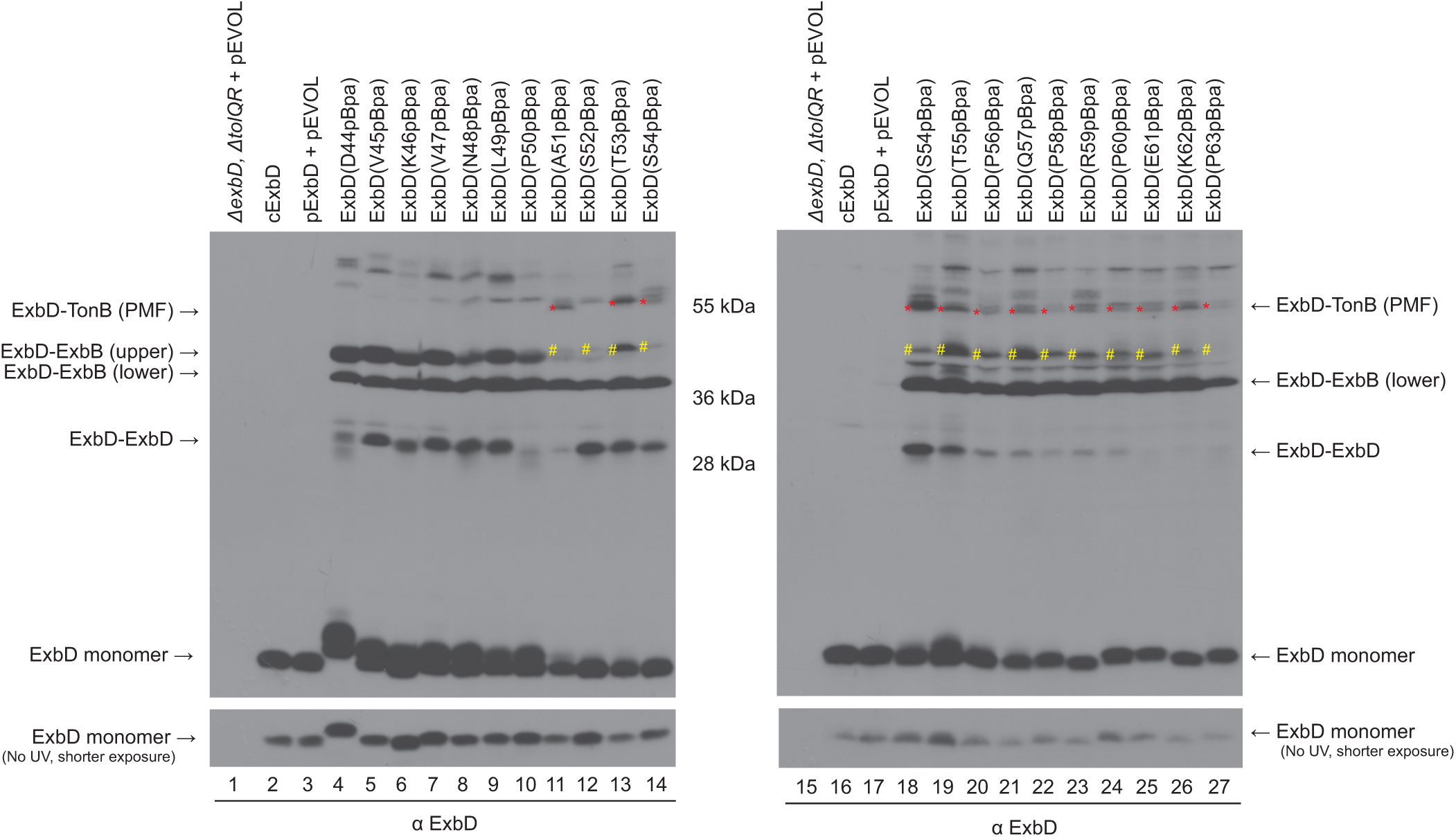
The majority of pBpa substitutions in the ExbD disordered region capture contacts with ExbB and ExbD. *In vivo* photo-cross-linking of ExbD pBpa substitutions in residues 44-54 (left panel) and 54-63 (right panel) from Fig. 4 probed with anti-ExbD antibodies. Note that ExbD (S54pBpa) is included on both immunoblots in lanes 14 and 18 for comparison of levels of expression. Strains were grown to mid-exponential phase, at which point photo-cross-linking was performed as described in Materials and Methods. Equivalent numbers of cells were precipitated with TCA and the ExbD-TonB complexes were visualized on immunoblots of 13% SDS-polyacrylamide gels with anti-ExbD polyclonal antibodies. cExbD designates chromosomally expressed ExbD from the wild-type strain, W3110. Plasmid pExbD (pKP999) and its corresponding ExbD pBpa substitutions were expressed in RA1045 (Δ*exbD*, Δ*tolQR*) with plasmid pEVOL. Positions of the PMF-dependent ExbD-TonB complex identified in Fig. 3, ExbD-ExbB (upper) and ExbD-ExbB (lower) from Figure 5A, the ExbD-ExbD from Figure 5B and ExbD monomer are indicated on the right and left. The ExbD-TonB (PMF) is identified by a red asterisk (*). The ExbD-TonB PMF-*in*dependent complex is distinguished from ExbB-ExbD (upper) by a yellow pound symbol (#). Mass standards are in the center. The steady-state ExbD levels from the same experiment were determined by immunoblot analysis of control samples that were not exposed to UV light (bottom). The plasmid identities and concentrations of propionate for chromosomal expression are listed in Table S1.

The ∼ 39 kDa ExbD-ExbB (lower) complex was formed by pBpa substitutions throughout the disordered domain from D44 through P63 at similar intensities. This indicates that a site in ExbD ∼20 residues from its predicted periplasmic TMD boundary [residues 32-43, (17)] binds to ExbB. The fact that ExbB has only a 20-residue periplasmic loop and a 21-residue periplasmic amino terminus raises the possibility that the ExbD disordered domain was condensed at some point during the TonB energization cycle such that it was closer to the inner-membrane (15). Alternatively, the ExbB periplasmic N-terminus might extend into the periplasm.

In contrast to the ExbD-TonB complexes, the ExbD-ExbD homodimers were captured more efficiently in the mutation-intolerant half of the ExbD disordered domain, residues 44-49 and 52-55, without emergence of a pattern indicative of structural elements.

### The ExbD disordered domain contains a conserved ΨXΨXLP motif

A multiple sequence alignment of the ExbD disordered domain was generated and displayed in a sequence logo using HHblits in the Gremelin.bakerlab program of all sequences within an expect score of e^-10^ of *Escherichia coli K-12 strain* (NP_417478.1) (Fig. 7A). The sequence logo identified a ΨXΨXLP motif where the psi symbol (Ψ) represents large aliphatic side-chained residues and X represents any non-hydrophobic residue (Fig. 7B). The motif consisted of residues V45, V47, L49, and P50, which reside in the mutation-intolerant segment of the disordered domain and were also conserved in the *Escherichia coli* ExbD paralog, TolR of the Tol system, but not in paralog MotB of the flagellar motor (Fig. 7B). There is modest crosstalk between the TonB and Tol systems such that they can substitute functionally for one another. However, to our knowledge, MotAB does not participate in any crosstalk. Cross-talk has been assumed to be due to the similarities between ExbB/ExbD TMDs of the TonB system and TolQ/TolR TMDs of the Tol system (21, 43, 50). Conservation of the ΨXΨXLP motif in both TolR and ExbD suggested that an additional source of cross-talk between ExbB/ExbD and TolQ/R might exist.

**Figure 7:**
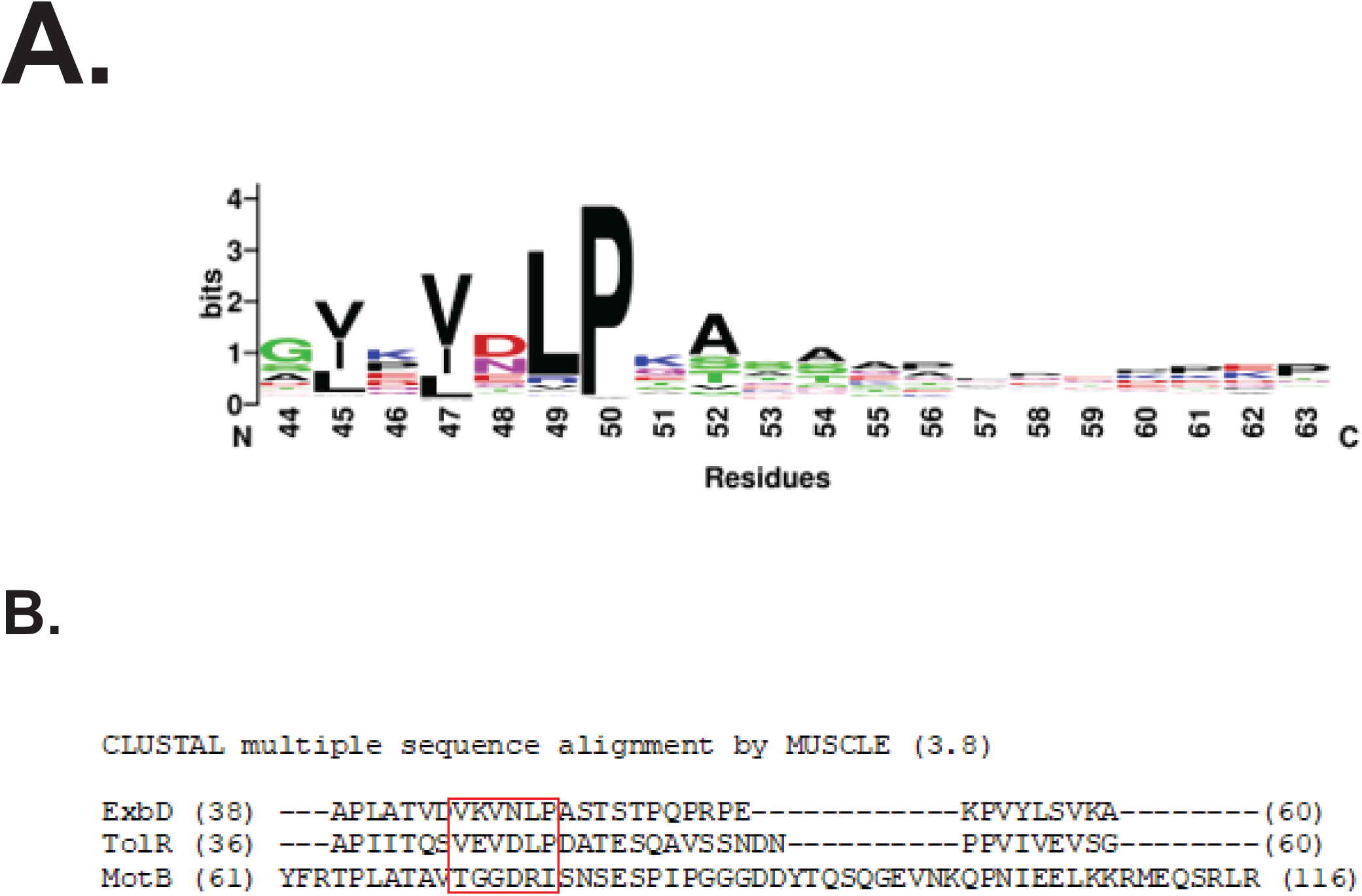
The ExbD disordered region contains conserved a ΨXΨXLP motif. The psi symbol (Ψ) represents a residue with a large aliphatic side chain and X represents any non-hydrophobic residue. (A) A WebLogo image of the ExbD disordered region. The image was generated from an ExbD multiple sequence alignment by HHblits of sequences within an E-value of e^-10^ of *Escherichia coli K-12* strain (NP_417478.1) using the Gremelin.bakerlab program (gremlin.bakerlab.org/). Positions with >75% sequences alignment gaps were removed. (B) A multiple sequence alignment was used to identify potential ΨXΨXLP motifs in ExbD homologs TolR and MotB. A clustal multiple sequence alignment was generated by MUSCLE (3.8) of *Escherichia coli* ExbD (AAA69172.1), TolR (PSM33938.1), and MotB (ACB03086.1). A red box encompasses the *Escherichia coli* K12 ExbD ΨXΨXLP motif containing conserved residues V45, V47, L49 and P50 and shows that these residues are conserved in TolR but not MotB.

### Only ExbD motif residues V45, V47, L49 and P50 are important for TonB system activity

Assessment of the activities of the pBpa substitutions in the mutation-intolerant segment showed that the least active ones [V45(pBpa) and V47(pBpa)], were part of the ΨXΨXLP motif (Fig. 4A). To better define the motif, individual Alanine substitutions were constructed in ExbD from residues 44 through 53. ExbD variants carrying V45A, V47A. L49A, and P50A mutations were the only ones with deficits in initial ^55^Fe-ferrichrome transport rates significantly below 90% of pExbD, with activities of 32%, 21%, 34%, and 50%, respectively (Fig. S3). Thus V45, V47, L49 and P50 were the only functionally important residues within the entire ExbD disordered domain.

To further define the motif, each of the V45A, V47A, L49A, and P50A substitutions in the motif region was combined with the others to create six double Ala substitutions. Had these mutations been simply additive in their effects, we would have seen that, for example, the V45A, P50A double mutant should have ∼16% activity (50% of 32%). Instead, this double mutant, like all six double mutants, was inactive in ^55^Fe-ferrichrome transport, suggesting a degree of synergy between them (Fig. 8A). However, the iron transport assays do not discriminate well among low levels of TonB system activity (51).

**Figure 8:**
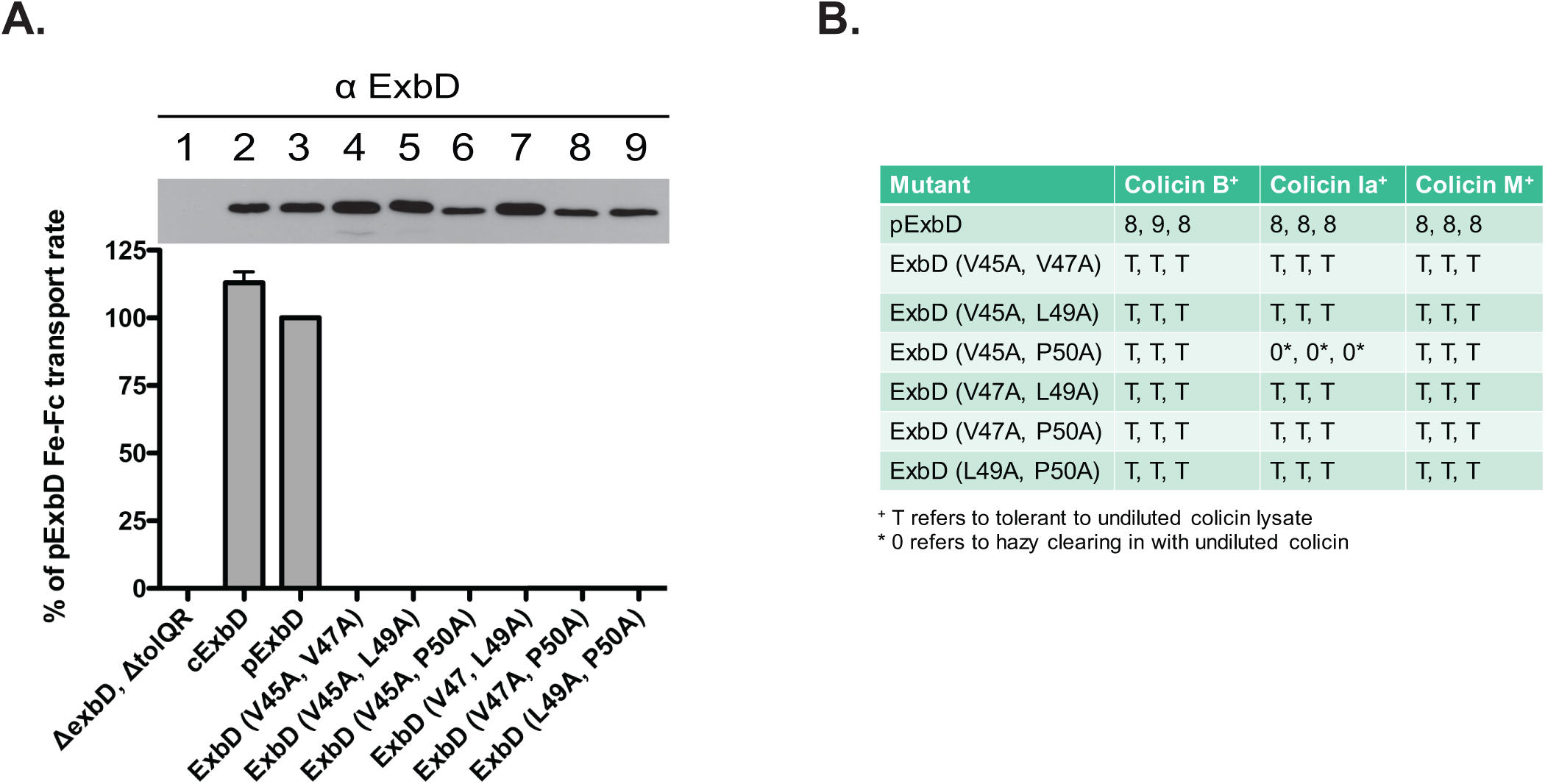
The synergy of double-Alanine substitutions within the ExbD ΨXΨXLP motif establishes its essentiality for TonB-dependent energy transduction. (A) Initial rate of ^55^Fe-ferrichrome transport for ExbD variants with the corresponding steady-state expression levels of ExbD proteins at the time of assay above each panel. Strains were grown to mid-exponential phase and assayed as described in Methods and Materials. The percentages of activity were calculated relative to the initial rate for ^55^Fe-ferrichrome transport of plasmid-encoded ExbD, pExbD (pKP999). Assays were performed in triplicate and carried out at least twice for each strain. The slopes were determined by a linear regression fit of scintillation counts per minute. The error bars indicate mean ± SEM for a representative triplicate set (above). Steady-state levels of ExbD-specific expression from the transport experiments are indicated above the corresponding bars of the graph below them. cExbD designates chromosomally expressed ExbD from the wild-type strain, W3110. The plasmid identities and concentrations of propionate for chromosomal expression are listed in Table S1. (B) Colicin sensitivity assay. Strains were grown to mid-exponential phase and assayed in triplicate with Colicin B, Colicin Ia, and Colicin M as described in the Material and Methods. pExbD (pKP999) and all ExbD Alanine substitutions were expressed from plasmids in RA1045 (Δ*exbD*, Δ*tolQR*). The plasmid identities and concentrations of propionate for chromosomal expression are listed in Table S1. Scoring was reported as the highest 5-fold colicin dilution that produced clearing in three different trials. “T” refers to colicin tolerance (insensitivity) and “0” refers to hazy clearing caused by the undiluted colicin lysate.

Colicins toxins are synthesized by some *E. coli* to kill sensitive strains. B-group colicins use the TonB system and its corresponding TBDTs to cross the outer membrane of *E. coli* to gain access to their targets in the periplasm, cytoplasmic membrane or cytoplasm (52). Colicin sensitivity assays, while not as discriminative among higher levels of TonB system activity, can detect the activity supported by as little as one molecule of TonB (51). To determine whether the double Ala substitutions could support any activity, their sensitivity to B-group colicins B, Ia, and M was tested (Fig. 8B). All of the double-Ala mutants were tolerant (insensitive) to virtually all colicins tested, suggesting that the motif residues do not work independently of each other and validating the identification of this motif.

### ExbD (V45A, V47A) and ExbD (V47A, L49A) stall TonB at Stage II of the energy transduction cycle

ExbD regulates TonB conformational changes as TonB moves through an energy transduction cycle (6, 34). Stage II of the TonB energization cycle is a PMF-*in*dependent Stage where the N-terminal ∼ 155 residues of TonB (an ∼23 kDa fragment) are protected from proteinase K degradation if full-length ExbD is present, although an actual complex between the two proteins had not been observed until this study (Fig. 5). Heretofore, Stage II had been detectable only when PMF was collapsed, preventing TonB from moving on to Stage III. When PMF, ExbB, ExbD, and TonB are all present and active, TonB automatically transitions from Stage II to Stage III, rendering TonB fully sensitive to proteinase K. Our observation that ExbD (44-48Ala) and ExbD (49-53Ala) failed to form a Stage III formaldehyde cross-link suggested that the effects of substitutions in the disordered domain motif might be to stall TonB at Stage II (Fig. S1). Since only ExbD residues V45, V47, L49, and P50 were important for ExbD activity, we assessed whether the inactive double mutants (V45A, V47A), (V47A, L49A) and (L49A, P50A) were required for TonB to proceed beyond Stage II based on its sensitivity to exogenously added proteinase K in spheroplasts. For comparison, we assessed the proteinase K accessibility of ExbD (D25N).

The hallmarks of the ExbD (D25N) mutant, where TonB is permanently stalled at Stage II are 1) that TonB forms a proteinase K-resistant fragment in both the presence and absence of CCCP and 2) that TonB is unable to remain in its stalled Stage II proteinase K-resistant conformation over time [Fig. 9, panel 4 +/- CCCP; (34)]. As expected, the Δ*exbD,* Δ*tolQR* strain did not support the ability of TonB to form the ∼23 kDa proteinase K-resistant TonB fragment [Fig. 9, panel 3]. A ∼28 kDa TonB fragment, seen previously and typified by Δ*exbD,* Δ*tolQR* and ExbD (D25N) in panels 3 and 4, does not appear to have relevance to the mechanism of TonB energization (34).

**Figure 9:**
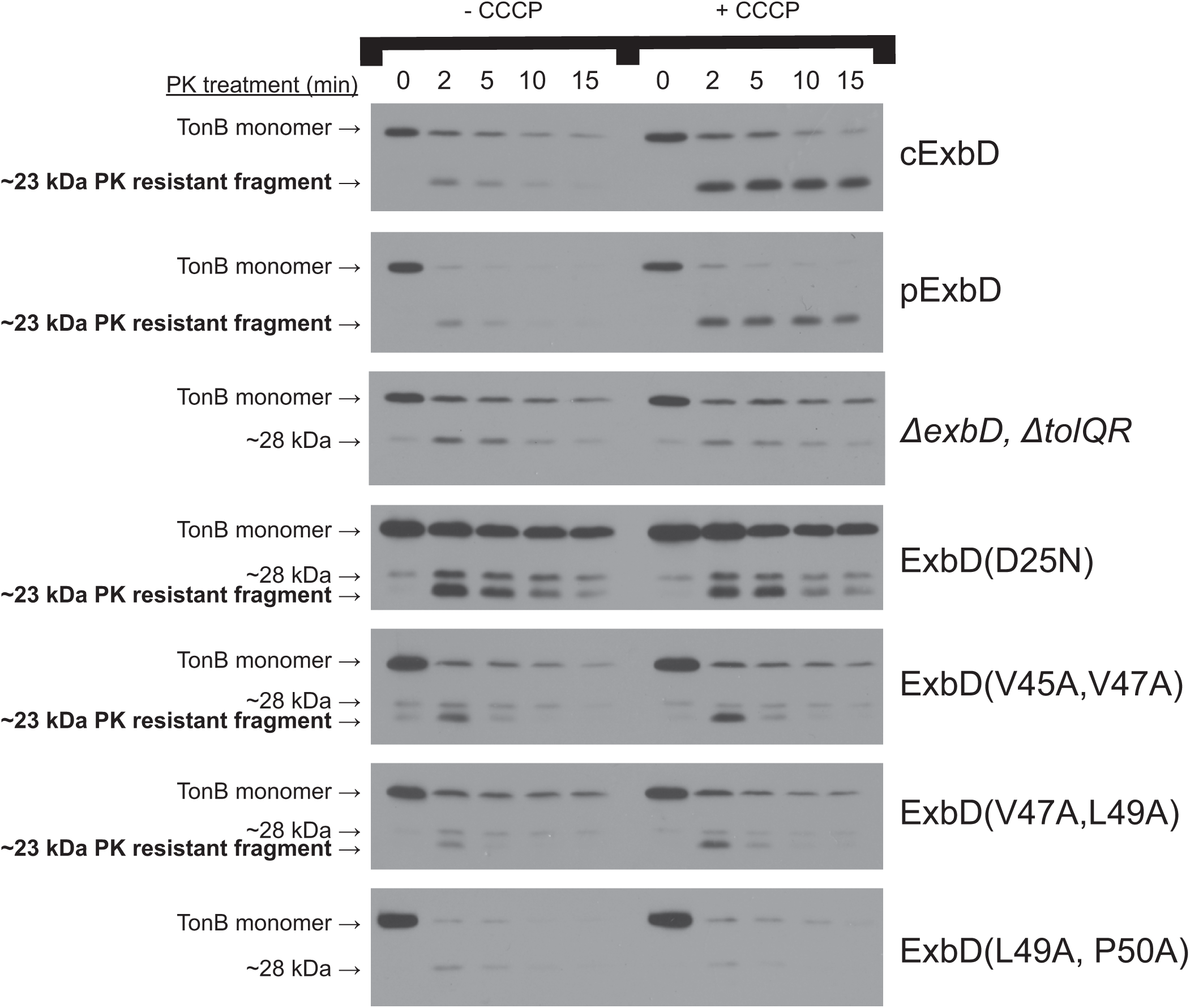
Mutations in the ExbD ΨXΨXLP motif leave TonB stalled at Stage II even when PMF is present. Spheroplasts were generated from W3110 (cExbD), and RA1045 (Δ*exbD*, Δ*tolQR*), as well as RA1045 containing plasmids pKP999 (pExbD), pKP1194 [ExbD (D25N)], pKP2171 [ExbD (V45A, V47A)], pKP2201 [ExbD (V47A, L49A)], and pKP2202 [ExbD (L49A, P50A)] as described in the Materials and Methods. Each spheroplast was treated with proteinase K (PK) for 0 min, 2 min, 5 min, 10 min and 15 min in the presence of DMSO (-CCCP) or CCCP dissolved in DMSO (+CCCP). The samples were visualized on immunoblots of 13% SDS-polyacrylamide gels with anti-TonB monoclonal antibodies. Identities of the various strains assayed are indicated on the left. Positions of TonB monomers, the ∼23 kDa PK-resistant TonB fragment indicative of Stage II of TonB energization and the ∼28 kDa fragment with no known relevance to any Stage in the TonB energization cycle are on the left (34). The appearance of the ∼23 kDa TonB fragment upon lengthy exposure of ExbD (L49A, P50A) samples allowed confident identification of the 28 kDa TonB fragment generated in that strain. The concentrations of propionate for chromosomal expression are listed in Table S1.

ExbD (V45A, V47A) and ExbD (V47A, L49A) largely mimicked the effect of ExbD (D25N) on TonB, leaving it stalled in Stage II (Fig. 9). Together the results suggest that the ExbD ΨXΨXLP motif was required for TonB to transition from Stage II to Stage III of the energization cycle. ExbD (L49A, P50A) was too sensitive to proteinase K to be evaluated (Fig. 9 panel 7).

### The ExbD disordered domain ΨXΨXLP motif is required for correct configuration of the ExbD C-terminus

To better understand why the ExbD ΨXΨXLP motif left TonB stalled in Stage II, we combined the V45A, V47A double mutation with pBpa substitutions along the length of ExbD: E14(pBpa)— presumably cytoplasmically localized based on the topology of ExbD; I24(pBpa)—in the ExbD TMD; S54(pBpa)—in the mutation-tolerant segment of the disordered domain; S67(pBpa)—just beyond the disordered domain; K97(pBpa)—a known point of ExbD-TonB contact through engineered Cys residues (39); and E136(pBpa)—near the C-terminus. With the exception of ExbD (I24pBpa), the other substitutions were active. Plasmid-encoded ExbD E14pBpa, S67pBpa, K97pBpa, and E136pBpa substitutions were 72%, 24%, 90%, and 100% of ^55^Fe-Ferrichrome transport of pKP999 + pEVOL in RA1045.

Overall, the double V45A, V47A substitutions did not have much effect on ExbD photo-cross-links to TonB through distal pBpa substitutions, except for (S54pBpa) where it greatly diminished the PMF-dependent ExbD-TonB complex (Fig. 10A lanes 6,7). ExbD (E14pBpa), in the cytoplasm, did not form complexes with TonB (Fig. 10A, lanes 4,5), however, ExbD (I24pBpa), in the TMD, formed a PMF-*in*dependent TonB complex (Fig. 10A lanes 6,7), indicating that at some point in its energy transduction cycle, TonB was in close proximity to ExbD through their TMDs. ExbD (K97pBpa) and ExbD (E136pBpa) also formed PMF-*in*dependent Stage II complexes with TonB (Fig. 10A, lanes 6,7,12-15). We do not know why the level of monomer was consistently reduced for ExbD (K97pBpa), but it clearly did not prevent capture of ExbD-TonB complexes.

**Figure 10:**
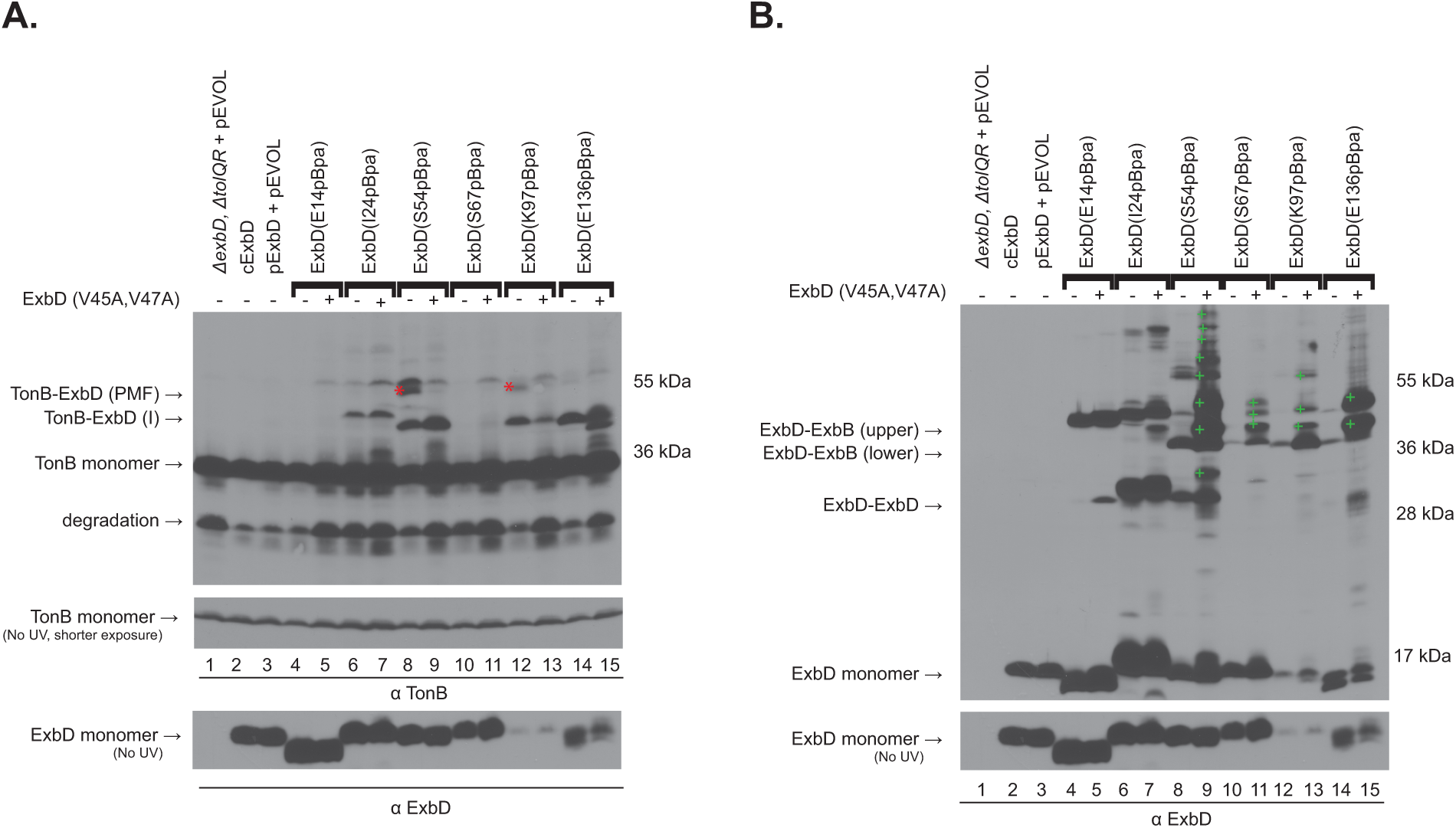
The N-terminal ExbD ΨXΨXLP motif is required for signal transduction to its C-terminus. *In vivo* photo-cross-linking of ExbD (E14pBpa), ExbD (I24pBpa), ExbD (S54pBpa), ExbD (S67pBpa), ExbD (K97pBpa), and ExbD (E136pBpa) with (+) and without (-) V45A, V47A double Alanine substitutions. Strains were grown to mid-exponential phase, at which point photo-cross-linking was performed as described in Materials and Methods. Equivalent numbers of cells were precipitated with TCA and the samples were visualized on immunoblots of 13% SDS-polyacrylamide gels with either (A) anti-TonB monoclonal antibodies or (B) anti-ExbD polyclonal antibodies. cExbD designates chromosomally expressed ExbD from the wild-type strain, W3110. The parent plasmid pExbD (pKP999) and corresponding ExbD variants were co-expressed with plasmid pEVOL in RA1045 (Δ*exbD*, Δ*tolQR*). (A) On the left, positions of a PMF-dependent ExbD-TonB complex [ExbD-TonB (PMF)—red asterisk], a PMF-*in*dependent complex [ExbD-TonB (I)], monomeric TonB, and a TonB degradation product are shown (See Fig. 3). Mass markers are indicated on the right. Steady state ExbD and TonB monomer levels from the same experiment were determined by immunoblot analysis of control samples that were not exposed to UV light (bottom). (B) On the left, positions of ExbD-ExbB (upper) and ExbD-ExbB (lower) (see Figure 5A), the ExbD-ExbD homodimer (see Figure 5B) and ExbD monomer are shown. Unidentified complexes trapped by the ExbD pBpa variants containing V45A,V47A are indicated on the immunoblot by a green (+) symbol. Mass markers are indicated on the right. Steady-state ExbD levels from the same experiment were determined by immunoblot analysis of control samples that were not exposed to UV light (bottom). The plasmid identities and concentrations of propionate for chromosomal expression are listed in Table S1.

Among the periplasmic pBpa substitutions tested, the presence of the V45A, V47A double substitution caused ExbD to photo-cross-link to a variety of unknown proteins, especially ExbD V45A, V47A (S54pBpa), which captured approximately eight different complexes with unknown proteins (Fig. 10B, lane 9). Furthermore, ExbD (E136pBpa) V45A, V47A captured two prominent complexes with unknown proteins (Fig. 10B, lane 15). These results showed that the conserved motif within the disordered domain of ExbD was important for correctly regulating contacts made by the ExbD C-terminus.

## DISCUSSION

TonB-dependent energy transduction is cyclic, with TonB, anchored in the inner membrane, engaging and disengaging from a TBDT as it energizes active transport across the outer membrane. Understanding the molecular interactions among TonB system proteins in the cytoplasmic membrane--ExbB, ExbD, TonB--and the TBDTs in the outer membrane is essential to understanding the mechanism of this novel dual-membrane transport system.

Our current model consists of multiple Stages that occur prior to productive TonB energy transmission to a TBDT [Fig. 11; (24, 34)]. In Stage I, TonB and ExbD homodimers are each found in separate complexes with ExbB homotetramers which do not interact. In Stage II, the ExbB-TonB and ExbB-ExbD oligomeric complexes come together to form a multimeric heterocomplex consisting of all three proteins, in which the first ∼155 residues of TonB are protected from digestion by exogenous addition of proteinase K to spheroplasts. Prior to our work here, Stage II could only be detected when the PMF was artificially collapsed by addition of the protonophore CCCP. We also knew that ExbD was somehow required to form the proteinase K-resistant fragment of TonB. However, prior to this study, we had not been able to detect the TonB-ExbD PMF-*in*dependent complex in Stage II, although its existence was implied. Nor had we been able to determine precisely through which residues the PMF-*in*dependent Stage II complex formed. From Stage II the TonB system transitions to Stage III, characterized by a requirement for PMF and a formaldehyde-cross-linked complex containing both ExbD and TonB. The final Stage in our current model is Stage IV in which TonB that has been properly reconfigured by ExbD binds the TBDT in such a manner that active transport across the OM occurs (6). After Stage IV, the multimeric heterocomplex of ExbB, ExbD, and TonB breaks down into isolated complexes of ExbB-ExbD, ExbB-TonB, and a minor fraction of free-floating ExbB (24)

**Figure 11:**
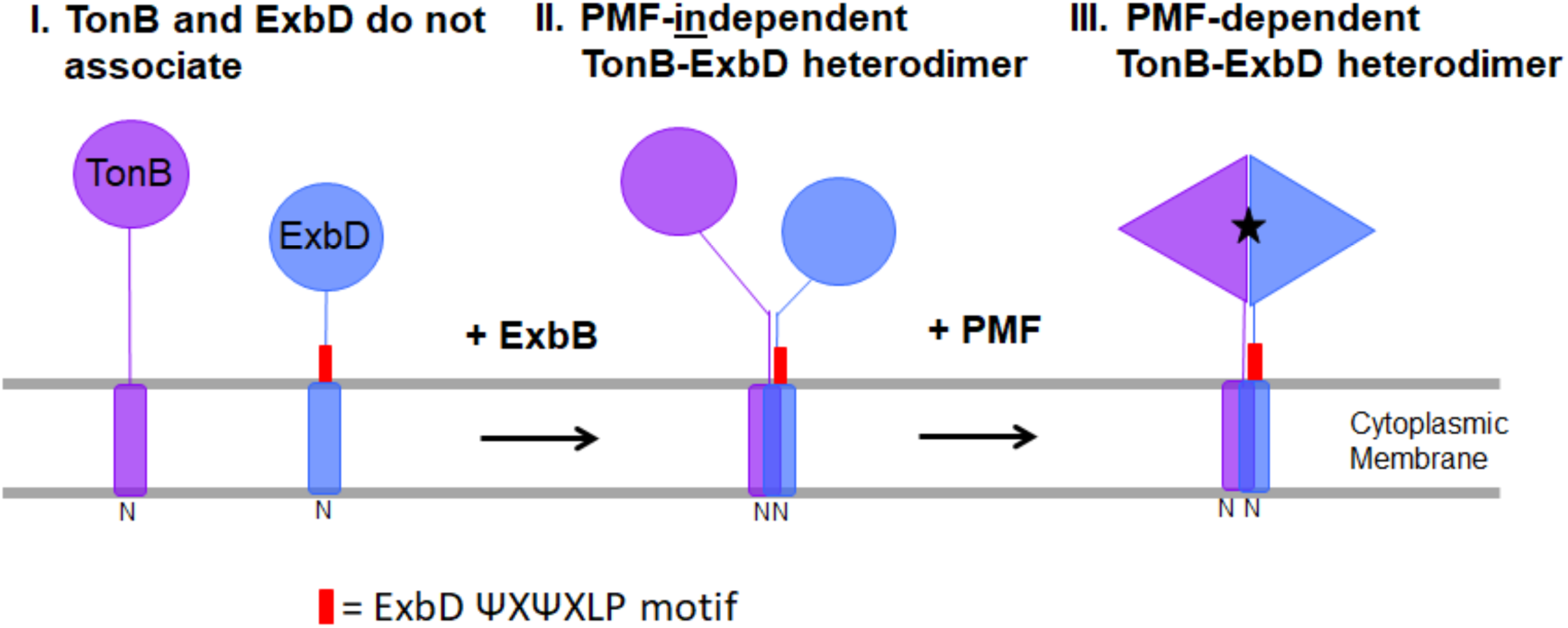
The ExbD ΨXΨXLP motif is required for TonB to respond to PMF in vivo. Model for the initial Stages of TonB energization [adapted from ref (24) with permission from the publisher]. This updated model covers the behavior of a single TonB. For an expanded model that accounts for behavior of TonB homodimers in the same context, see (24)]. In Stage I, ExbD and TonB do not interact, although both are stabilized by ExbB homotetramers (not shown). In Stage II, the ExbD TMD, adjacent ΨXΨXLP motif (red box), and the rest of the ExbD disordered domain directly interact with TonB in a PMF-*in*dependent manner, indicating their in vivo proximity. The C-termini of ExbD and TonB do not interact. In Stage III, the conformational relationship between ExbD and TonB changes in response to PMF such that TonB is correctly configured by ExbD for energy transmission to a TBDT, a transition for which the ExbD ΨXΨXLP motif is required. Stage IV (not shown) is interaction of TonB with a TBDT. TonB is then recycled back to Stage I.

### The ExbD disordered domain is highly dynamic

*In vivo* photo-cross-linking has the advantage of potentially capturing every interaction within ∼3 angstroms that a single pBpa substitution is involved in during the TonB energization cycle (46). Throughout the ExbD disordered domain (residues 44-63 immediately distal to its TMD), each single ExbD residue captured multiple complexes. Disordered domains of proteins are known to have a high degree of conformational plasticity that can mediate changes in protein oligomerization and function (53). For example, substrate binding to the intrinsically disordered domain of periplasmic protease/chaperone DegP causes a change from proteolytically inactive hexamers to proteolytically active polyhedrons consisting of 4, 6, and,8 trimers (54, 55).

Consistent with characteristics of disordered domains, most of the ExbD pBpa substitutions within the disordered domain captured at least four out of five identifiable complexes (ExbD homodimer, two ExbD-ExbB complexes, and two ExbD-TonB complexes) through a single pBpa substitution. This indicated that residues within the ExbD disordered domain underwent multiple partner swaps during the Stages of the TonB energization cycle and quantified the minimum number of disordered domain contacts that occurred.

#### ExbD-TonB complexes

TonB is predicted to have large regions of intrinsic disorder starting immediately after its TMD, suggestive of multiple sites through which protein-protein interactions occur (56). Consistent with that, the dimeric crystal structures of the TonB C-terminus do not exist in vivo, even though TonB homodimerizes through that domain in vivo (26). Because TonB contains no single essential residue (26, 36, 56–60), ExbD is hypothesized to determine the final configuration of TonB that is capable of transducing energy to TBDTs (21, 26, 29, 34). As pBpa substitutions, different subsets of residues within the ExbD disordered domain (residues 44-63) captured both PMF-*in*dependent and PMF-dependent ExbD-TonB interactions using *in vivo* photo-cross-linking. For the first time, a PMF-*in*dependent ExbD-TonB complex was captured, which validated the previously predicted existence of Stage II in the TonB energization cycle.

A PMF-*in*dependent Stage II complex was formed by residues from 44-63--nearly all the disordered domain. The size of the ExbD disordered domain that captured this interaction was much smaller than the ∼ 155 residues of TonB protected against proteinase K degradation by ExbD (6). Either the full length of ExbD (∼ 141 residues) can directly protect TonB or, as has been proposed previously, the disordered domain of ExbD induces conformational changes that render the TonB protease-insensitive (21, 60). Previous disulfide cross-linking experiments with engineered Cys substitutions within ExbD residues 92-121 failed to capture a PMF-*in*dependent interaction (38, 39).

PMF-dependent ExbD-TonB complexes were captured only by the more distal end of the disordered region through ExbD residues A51, and T53 to P63. We previously showed that deletions of residues 42-51 and 52-61 prevented formation of the PMF-dependent Stage III ExbD-TonB complex, but did not know if those regions were directly or indirectly responsible (35). We showed here that the ExbD disordered domain directly contacted TonB PMF-dependently. Thus, we identified another ExbD region besides residues 92-121 that requires PMF for interaction with TonB. However, like the ExbD region of 92-121 Cys substitutions, Ala substitutions for the ExbD disordered domain residues that captured the PMF-dependent ExbD-TonB interaction did not significantly decrease TonB system activity. This may suggest that the disordered domain and the 92-121 residue region are simply the read-outs for PMF-dependent events initiated elsewhere in ExbD, possibly through its TMD, which contains the only potentially PMF-responsive residue in the TonB system (15, 29, 36). It also cannot be ruled out that some functionally important ExbD disordered residues (i.e. V45 and V47) directly interact with TonB PMF-dependently, but due to their low activity, the interaction was not captured.

#### ExbD-ExbB complexes

ExbD and ExbB formaldehyde cross-link into a complex of complex of ∼41 kDa (29).Surprisingly, two distinct ExbD-ExbB complexes were captured via photo-cross-linking: an ∼ 39 kDa (lower) complex and an ∼43 kDa (upper) complex. While both techniques are probes for close contacts, formaldehyde cross-linking occurs through residues W, H,Y,R,C, K and N-termini and requires that both proteins have these specific residues in close proximity for the cross-link to occur (48). Photo-cross-linking, on the other hand, allows crosslinking to occur between the engineered site containing pBpa and any carbon-hydrogen bonds within ∼ 3 angstroms (46). In addition, the apparent mass of a complex can vary with the sites through which the two proteins are linked (61). The difference in the masses of formaldehyde cross-linked and photo-cross-linked may therefore be due to differences in the sites of complex formation. We can say, however, at a minimum, there are two different forms ExbD-ExbB complexes that suggest a single pBpa residue binds to two different regions of ExbB as the energy transduction cycle proceeds. If so then, it would suggest that in only one of these interactions are the formaldehyde-cross-linkable residues in correct apposition in both ExbD and ExbB.

The ∼ 39 kDa ExbB-ExbD (lower) complex was captured with similar intensities by all residues in the ExbD disordered domain. ExbB has few of its residues in the periplasm: an ∼ 21-residues N-terminus and ∼ 26-residue loop between TMDs 2 and 3 (14, 15). The topological relationship between the ExbD disordered domain and ExbB suggest that either ExbB sequences assume extended conformations or that the ExbD disordered domain moves close to the membrane at some point during the TonB energization cycle. The latter possibility would be consistent with one of the three ExbB-ExbD structures isolated and mapped via 3D electron microscopy with ratios 4:2 in which electron density of the ExbD periplasmic domain is parallel with the membrane (62). It would also be consistent with a genetic suppressor of mutants in the ExbB paralog TolQ that map to C-terminus of the ExbD paralog, TolR in the Tol system (63).

The ∼43 kDa ExbD-ExbB (upper) complex was captured essentially only through the six residues immediately distal to ExbD TMD. Unlike the ∼ 39 kDa ExbD-ExbB complex described above, this one could be based solely on proximity of the reactive ExbD pBpa residue to periplasmic ExbB residues that were equally adjacent to the cytoplasmic membrane. Taken together, the difference in apparent masses and different interaction profiles of the ∼ 39 and ∼ 43 kDa complexes indicated that ExbD was in two different conformations in each complex and likely interacting with different parts of ExbB.

When the TonB C-terminus is intact, its extreme N-terminus (L3C) is protected from labeling by the thiol-specific membrane-permeant reagent Oregon Green^®^ 488 maleimide; when the TonB C-terminus is absent, TonB L3C is labeled (58). This behavior suggests two different conformations of ExbB cytoplasmic domains, one which protects the TonB L3C from labeling during binding of TonB to a TBDT and one which allows TonB L3C labeling when the TonB C-terminus is engaged with ExbD (28). Perhaps, the two photo-cross-linked ExbD-ExbB complexes in this work are related to ExbB conformational changes that influence the Oregon Green^®^ 488 maleimide labeling of cytoplasmic tail mutant TonB (L3C) as TonB moves through an energy transduction cycle. Addition of CCCP did not prevent formation of the ExbB-ExbD (upper) or ExbB-ExbD (lower) photo-cross-linked complexes, ruling out the possibility that either was dependent on PMF.

#### ExbD-ExbD complexes

An ExbD homodimer (∼31 kDa) identified using *in vivo* photo-cross-linking had a similar apparent mass as the formaldehyde-cross-linked ExbD homodimer (∼30 kDa) (29). Except for ExbD (P50pBpa) and ExbD (A51pBpa), the intensity of the ExbD homodimer complex gradually decreased as residues grew more distant from the TMD. Because the residues that capture ExbD homodimers are immediately distal to its TMD, it suggested that the ExbD TMDs were also homodimerized.

### The *in vivo* data compared to recent ExbB/ExbD structures

The ratio of cytoplasmic membrane proteins ExbB, ExbD and TonB in vivo is 7ExbB:2ExbD:1TonB under four different conditions of iron availability (13). This oddly asymmetric ratio played an important role in our model of the TonB system (24, 30). In vitro, the ratios of these proteins in purified structures are varied with none of them actually accounting for the cellular ratio, and with TonB protein included in only one: 4ExbB: 2ExbD and 4ExbB: 1ExbD: 1TonB from single particle EM studies (49, 62); 5ExbB:1ExbD and 6ExbB:3ExbD from crystal structure determinations (64, 65); 6ExbB:1ExbD from mass spectrometry (66); and most recently, 5ExbB: 2ExbD from cryo-EM (67).

Recently, two crystal structures and a cryo-EM structure provide the first detailed view of ExbB monomers, arranged front to back and forming a pore: 5 ExbB with 1 ExbD TMD in the pore (crystal); 6 ExbB: 3 ExbD TMD in the pore (crystal); 5 ExbB: 2 ExbD TMD in the pore (cryo-EM) (64, 65, 67). The ExbB monomers are a major and very important accomplishment and are consistent with the known topology of ExbB; the oligomeric structures diverge from what is known *in vivo* about ExbB oligomers, ExbD and TonB:

1. In vivo, ExbB formaldehyde-crosslinks into tetramers that are dimers of homodimers. ExbB can also form disulfide-linked homodimers through D211C (30). The D211 residues are not sufficiently close in the front-to-back configuration of ExbB monomers in crystal and cryo-EM structures to form disulfide-linked homodimers (64, 65, 67).
2. In vivo, ExbB formaldehyde-cross-links to TonB and TonB interacts with ExbB TMD1 (15, 29, 30, 68). None of the recent ExbB crystal or cryo-EM pentameric/hexameric structures includes TonB (64, 65, 67).
3. In the recent ExbB crystal and cryo-EM structures, TonB is proposed to be outside the pore formed by the ExbB pentamer/hexamers because the lumens of the pores are entirely filled with one-to-three ExbD TMDs (64, 65, 67). Because TonB is known to go through its energy transduction cycle homodimerized through or near its TMD (24), the crystal and cryo-EM structures give rise to a ratio of 5-6 ExbB:10-12 TonB—the converse of the cellular ratio of 14 ExbB:2 TonB.
4. The *in vivo* photo-cross-linking data from this study indicated that ExbD (I24pBpa) in the ExbD TMD, as well as most of the immediately adjacent periplasmic ExbD disordered domain captured the ExbD-TonB PMF-*in*dependent complex. This strongly suggested that ExbD and TonB TMDs interact in vivo. As noted in point 3 above, neither the recent crystal nor cryo-EM structures of ExbB oligomers permits ExbD and TonB to interact through or near their transmembrane domains because the lumens of the ExbB pores are fully occupied by ExbD TMDs (64, 65, 67).

The discrepancies between *in vivo* results and ExbB crystal and cryo-EM structures raise questions about the validity of the structures and their possible roles in TonB-dependent energy transduction. Our current model identifies Stages in that process (24), and in this study, we identified dynamic contacts among ExbD, TonB and ExbB that occurred during those Stages through identical pBpa substitutions, changing as TonB moved through an energy transduction cycle. Perhaps the *in vitro* structures approximate a version of Stage I where the ExbB tetramers bind ExbD homodimers or TonB homodimers, but not both together (24).

### The ExbD ΨXΨXLP motif is required for signal transduction

Single Ala substitutions in the ExbD ΨXΨXLP motif (V45, V47, L49, P50) decreased but did not eliminate TonB system activity. Evidence for its importance as a motif came from combining the Ala substitutions, all of which double combinations were virtually inactive, even in the most sensitive assays capable of detecting one active molecule of TonB (51).

In this study we discovered that the ExbD ΨXΨXLP motif (V45, V47, L49, P50) played a role in signal transduction to both TonB and to its own C-terminal domain. First, we were able to identify the step mediated by the ΨXΨXLP motif within the context of our current model for TonB energization (Fig. 11). The double mutation (V45A, V47A) prevented formation of Stage III PMF-dependent ExbD-TonB complexes mediated through S54pBpa and K97pBpa, stalling the TonB energization cycle in Stage II, just after ExbD and TonB were brought together, but before they were able to form the PMF-dependent complex. Consistent with that, the double mutation (V45A, V47A) did not prevent formation of the PMF-*in*dependent ExbD-TonB complex, which ruled out the motif out as the site of contact.

Second, the V45A,V47A double mutation also caused pBpa substitutions in all four C-terminal ExbD residues tested (S54, S67, K97, and E136) to form complexes with unknown proteins. This was most notable in the case of S54pBpa where approximately eight new complexes with unknown proteins were captured. It appeared therefore that the ΨXΨXLP motif was important for transducing conformational signals to the ExbD C-terminus, which, when not properly directed, had a pluripotent capacity to interact widely.

## MATERIALS and METHODS

### Strains and plasmids

Strains and plasmids used in these studies are listed in Table S1. Strain KP1526 was constructed by P1*vir* transduction of Δ*exbD*::*cam* from RA1021 (W3110, Δ*exbD*::*cam*) into KP1525 (W3110, Δ*tolQR*, *aroB*) [RA1021 was a kind gift from Ray Larsen (35).] Plasmids encoding Alanine and amber substitution mutations in *exbD* were primarily pKP999 derivatives. Amber substitutions in plasmids encoding ExbD (V45A, V47A) were derived from pKP2171. To determine the identity of potential ExbB complexes, a His_6_ epitope tag was added to the C-terminus of ExbB in the plasmid pKP1657, which encodes the ExbBD operon with and without the ExbD D44_am_ mutation. Mutant plasmids were engineered using extralong PCR. Primers are available upon request. All genes containing PCR-based changes were verified by DNA sequencing at the Penn State Genomics Core Facility, University Park, PA.

### Media and growth conditions

Cultures were grown at 37°C with aeration. Saturated cultures in Luria-Bertani broth were sub-cultured in mid-exponential phase at 1:75 in M9 minimal media supplemented with 0.4% glycerol, 0.2% Casamino Acids, 40 µg/ml tryptophan, 4 µg/ml vitamin B1, 1 mM MgSO_4_, 0.5 mM CaCl_2_, and 37 µM FeCl_3_. Plasmid-encoded ExbD and its variants were expressed in RA1045 (W3110, Δ*exbD*, Δ*tolQR*) in the presence of 100 µg/mL ampicillin unless otherwise noted. Sub-cultures in M9 medium were induced with sodium propionate (ranging from 0-45 mM) to approximate the steady state levels of ExbD protein from W3110, the wildtype control (Table S1). Subcultures were harvested by immediate precipitation with ice-cold trichloroacetic acid (TCA) at an A_550_ ranging from 0.40 to 0.60 (40).

### *In vivo* formaldehyde-cross-linking

Equal numbers of bacteria were resuspended in sodium phosphate buffer (pH 6.8) and treated for 15 min at room temperature with 1% formaldehyde (Electron Microscopy Sciences Cat #15710) as previously described (29, 40). Following formaldehyde treatment, bacteria were suspended in sample buffer, heated to 60° C, and equal cell numbers were loaded in each well of 13% sodium dodecyl sulfate (SDS)-polyacrylamide gels (69). ExbD complexes and TonB complexes were detected by immunoblotting with ExbD-specific polyclonal antibodies and TonB-specific monoclonal antibodies, respectively (13, 70). Equal loads were confirmed by Coomassie staining of the PVDF membrane used for the immunoblot.

### *In vivo* photo-cross-linking

Plasmid pEVOL encodes an orthogonal tRNA synthetase and a corresponding orthogonal tRNA_CUA_ that recognizes amber (UAG) codons (45). Plasmids with amber substitutions in *exbD* were co-expressed with pEVOL in RA1045 (W3110, Δ*exbD*, Δ*tolQR*) and processed as previously described (7). At A_550_ of ∼0.2, sub-cultures in M9 were treated with 0.004**%** arabinose to induce expression of the tRNA synthetase/tRNA pair from pEVOL, 0.8 mM the photo-cross-linkable amino acid p-Benzoyl-L-phenylalanine (pBpa; Bachem AG, www.bachem.com) and levels of propionate required for ExbD protein expression at near-chromosomal levels (Table S1). 2.0 A_550_-ml were harvested at A_550_ ∼ 0.5, pelleted at room temperature and resuspended in 2 mL supplemented M9 media (as described above with 1.85 µM FeCl_3_ instead of 37 µM, and without antibiotics, pBpa, and propionate). Equal volumes of each sample were transferred into single wells of each of two 24-well plates (NEST Cell Culture Plate, cat: 702001). One plate was irradiated with 365-nm light for 30 min on ice and the second plate was incubated in the dark simultaneously for 30 min on ice. Samples were precipitated with 10% TCA as described previously and processed for immunoblot analysis with ExbD-specific polyclonal antibodies and TonB-specific monoclonal antibodies (13, 70). Equal loads were confirmed by Coomassie staining of the PVDF membrane used for the immunoblot.

### Initial rates of ^55^Fe-ferrichrome and ^55^Fe-enterochelin transport

Cultures were grown as described in media and growth conditions. Strains harboring plasmids with and without amber substitutions in *exbD* were grown as described in the *in vivo* photo-cross-linking section. ^55^Fe-ferrichrome assay and ^55^Fe-enterochelin assay were performed in triplicate as previously described (40). To exclude competition from native enterochelin, ^55^Fe-enterochelin uptake was assessed in *aroB* strains that were sub-cultured with additional phenylalanine (40 μg/ml) and tyrosine (40 μg/ml) to compensate for their inability to synthesize aromatic amino acids (7). To determine steady state levels of the different ExbD variants at the time of assay, samples were precipitated with 10% TCA at the start of the assay and equivalent numbers of cells processed for immunoblot analysis on 13% SDS-polyacrylamide gels. ExbD proteins were detected by immunoblotting with ExbD-specific polyclonal antibodies (13, 70).

### Ethidium bromide accumulation

Sub-cultures were grown in M9 medium as described above. At *A*_550_ of 0.5, 2 mL of cells were transferred to 4.5 mL disposable cuvettes (VWR Cat# 30620-276). Ethidium bromide (6.25 μM) was added to the cells, which were excited at 520 nm and fluorescence was detected using a Perkin-Elmer LS-55 fluorescence spectrophotometer at 590 nm. The width of the excitation and emission slits was 15 nm. Fluorescence emission data were collected every second for 3 minutes using the Time Drive application from the FLWINLAB software. For CCCP controls, either 15 μM or 60 μM CCCP was added two minutes prior to ethidium bromide. The ethidium bromide accumulation rate for each sample was determined by linear regression and recorded as Fluorescence Units per second (FU/sec) (42).

### Proteinase K Accessibility

TonB sensitivity to proteinase K was performed as previously described (35). Briefly, cultures were grown as described in M9 minimal media and converted to spheroplasts. At 5 min prior to proteinase K treatment (25 μg/mL), samples were treated with 60 µM CCCP dissolved in dimethylsulfoxide (DMSO), where indicated. Samples were collected at 2 min, 5 min, 10 min, and 15 min following proteinase K treatment with the addition of phenylmethylsulfonyl fluoride (1.0 mM) to inactivate proteinase K at the specified times. Samples were precipitated with 10% TCA final concentration, suspended in sample buffer, loaded with equal volumes on 13% SDS-polyacrylamide gels and detected via immunoblotting with TonB-specific monoclonal antibodies (13, 70).

### Colicin Sensitivity Assay

The colicin sensitivity assay was performed as previously described (40). Strains expressing ExbD at chromosomal levels were grown to mid-exponential phase in T-broth with 100 µg/ml ampicillin and with levels of propionate that induced chromosomal levels of expression as confirmed by TCA precipitation of cultures and comparison to W3110 on immunoblots. Strains were plated in T-top agar on T-plates containing the same propionate concentrations and ampicillin as were used in the sub-cultures. Five-fold dilutions of Colicins B, Ia, or M (Colicin B enters via FepA; Colicin Ia enters via CirA; Colicin M enters FhuA) were spotted on the bacterial lawns and were incubated overnight at 37°C. Scoring was reported as the highest dilution that produced clearing. Sensitivity to each plasmid was assayed in triplicate.

## ACKNOWLEDGEMENTS

Support from NIAID for grant AI11322 to K.P. is gratefully acknowledged. We thank Erica Johnson, Dakota Matson, and Molly Sweeney for engineering many of the mutant plasmids in this study. We thank Anne Ollis for construction of strain KP1526.

## SUPPLEMENTAL INFORMATION

**Table S1.** Kopp and Postle:

| Strain | Genotype |  | Reference |
| --- | --- | --- | --- |
| W3110 | F− IN(*rrnD-rrnE*)1 |  | [1] |
| KP1270 | W3110, *aroB* |  | [2] |
| KP1509 | W3110 Δ*exbD* Δ*tolQR*, Δ*tonB::kan* | | [5] |
| KP1526 | W3110, Δ*exbD*, Δ*tolQR*, *aroB* | | Present Study |
| RA1017 | W3110, Δ*exbBD::kan*, Δ*tolQRA* | | [4] |
| RA1045 | W3110, Δ*exbD*, Δ*tolQR* | | [3] |
| Plasmid | Genotype or Phenotype | <sup>a</sup>Propionate Induction Concentration | References |
| pEVOL-BpF | tRNA synthetase and tRNA for pBpa insertion, *cam*<sup>r</sup> | N/A | [6] |
| pKP999 | *exbD* in pPro24, *amp*<sup>r</sup> | 0.5 mM | [5] |
| pKP1657 | *exbBD* in pPro24, *amp*<sup>r</sup> | 0 mM | Present study |
| pKP1064 | ExbD (D25N) | 0 mM | [5] |
| pKP2073 | ExbD (44-48Ala) | 0.25 mM | Present study |
| pKP2077 | ExbD (49-53Ala) | 0.5 mM | Present study |
| pKP2153 | ExbD (54-58Ala) | 0.75 mM | Present study |
| pKP2180 | ExbD (59-63Ala) | 20 mM | Present study |
| pKP2140 | ExbD (D44A) | 0.25 mM | Present study |
| pKP2141 | ExbD (V45A) | 2.5 mM | Present study |
| pKP2142 | ExbD (K46A) | 0 mM | Present study |
| pKP2143 | ExbD (V47A) | 1 mM | Present study |
| pKP2144 | ExbD (N48A) | 0.25 mM | Present study |
| pKP2145 | ExbD (L49A) | 0.25 mM | Present study |
| pKP2154 | ExbD (P50A) | 0 mM | Present study |
| pKP2152 | ExbD (S52A) | 0.25 mM | Present study |
| pKP2146 | ExbD (T53A) | 0.25 mM | Present study |
| pKP2007 | ExbD (D44Am) | 0 mM | Present study |
| pKP2133 | ExbD (V45Am) | 0 mM | Present study |
| pKP2017 | ExbD (K46Am) | 0 mM | Present study |
| pKP2131 | ExbD (V47Am) | 0.5 mM | Present study |
| pKP2018 | ExbD (N48Am) | 2 mM | Present study |
| pKP2147 | ExbD (L49Am) | 0.5 mM | Present study |
| pKP1971 | ExbD (P50Am) | 0 mM | Present study |
| pKP2138 | ExbD (A51Am) | 45 mM | Present study |
| pKP1951 | ExbD (S52Am) | 1 mM | Present study |
| pKP2151 | ExbD (T53Am) | 1 mM | Present study |
| pKP1972 | ExbD (S54Am) | 1 mM | Present study |
| pKP2139 | ExbD (T55Am) | 8 mM | Present study |
| pKP1973 | ExbD (P56Am) | 5 mM | Present study |
| pKP2137 | ExbD (Q57Am) | 10 mM | Present study |
| pKP1974 | ExbD (P58Am) | 5 mM | Present study |
| pKP1898 | ExbD (R59Am) | 5 mM | Present study |
| pKP1975 | ExbD (P60Am) | 8 mM | Present study |
| pKP2027 | ExbD (E61Am) | 10 mM | Present study |
| pKP1949 | ExbD (K62Am) | 8 mM | Present study |
| pKP2136 | ExbD (P63Am) | 7 mM | Present study |
| pKP1926 | ExbD (E14Am) | 4 mM | Present study |
| pKP2064 | ExbD (I24Am) | 0 mM | Present study |
| pKP2167 | ExbD (S67Am) | 5 mM | Present study |
| pKP2227 | ExbD (K97Am) | 2 mM | Present study |
| pKP1907 | ExbD (E136Am) | 0 mM | Present study |
| pKP2171 | ExbD (V45A, V47A) | 1 mM | Present study |
| pKP2164 | ExbD (V45A, L49A) | 0.5 mM | Present study |
| pKP2182 | ExbD (V45A, P50A) | 0.25 mM | Present study |
| pKP2201 | ExbD (V47A, L49A) | 0.25 mM | Present study |
| pKP2190 | ExbD (V47A, P50A) | 0.25 mM | Present study |
| pKP2202 | ExbD (L49A, P50A) | 0.5 mM | Present study |
| pKP2213 | ExbD (V45A,V47A, E14Am) | 0 mM | Present study |
| pKP2214 | ExbD (V45A,V47A, I24Am) | 0 mM | Present study |
| pKP2161 | ExbD (V45A,V47A, S54Am) | 3 mM | Present study |
| pKP2215 | ExbD (V45A,V47A, S67Am) | 1.5 mM | Present study |
| pKP2228 | ExbD (V45A,V47A, K97Am) | 30 mM | Present study |
| pKP2220 | ExbD (V45A,V47A, E136Am) | 0 mM | Present study |
| pKP2063 | ExbD (S54pBpa, D25N) | 1 mM | Present study |
| pKP1932 | ExbD (R59pBpa, D25N) | 5 mM | Present study |
| pKP2259 | ExbD (S54AM)-His(6x) | 5mM | Present study |
| pKP2295 | ExbB-ExbD (D44Am) | 0 mM | Present study |
| pKP2272 | ExbB-His(6x)-ExbD (D44Am) | 0 mM | Present study |

<sup>a</sup> Propionate concentration required for plasmid induction in RA1045 background strain to achieve similar W3110 ExbD chromosomal protein levels.

## SUPPLEMENTAL FIGURES Kopp and Postle

**Figure S1:**
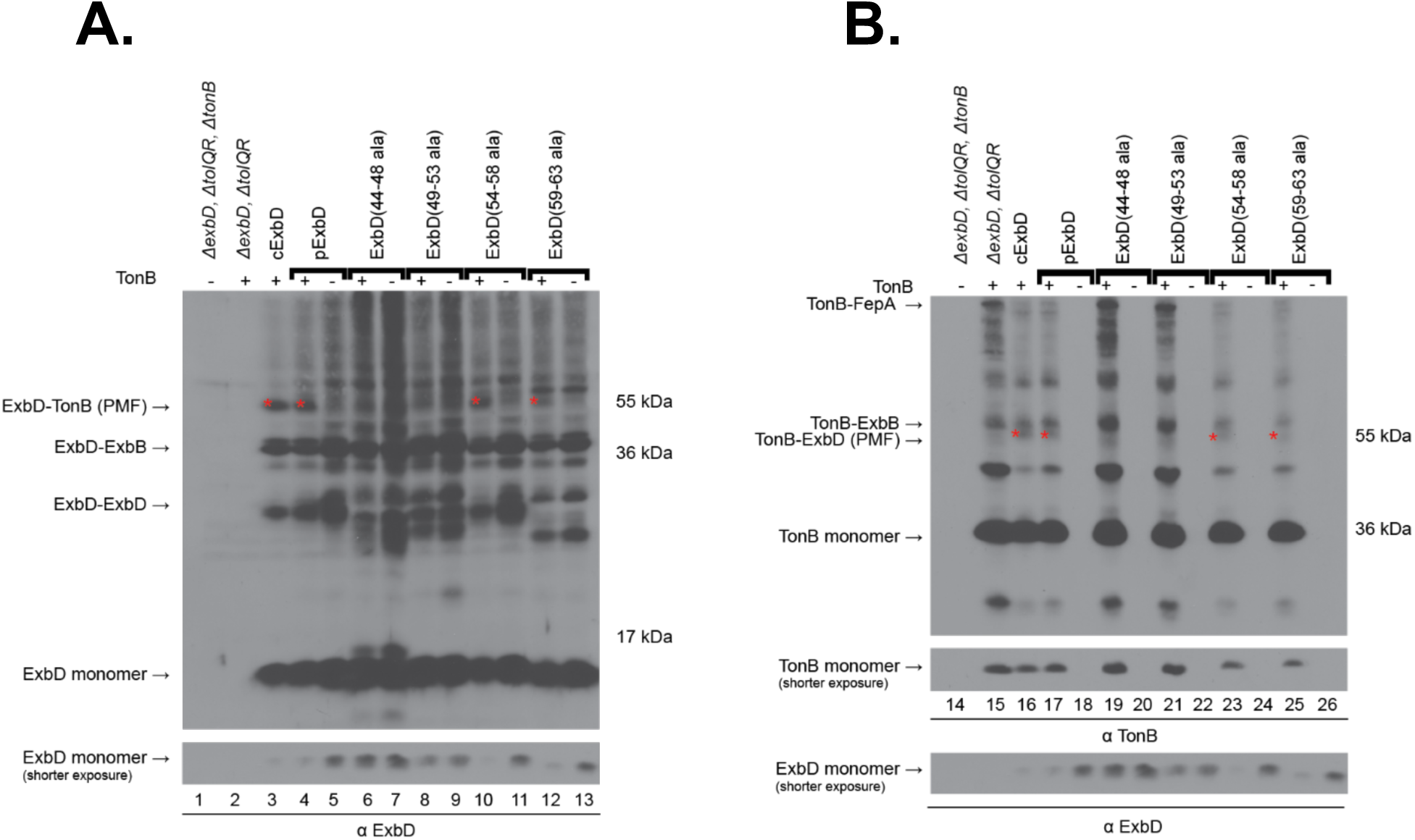
ExbD (44-48Ala) and ExbD (49-53Ala) fail to form a formaldehyde crosslinked ExbD-TonB PMF-dependent complex. *In vivo* formaldehyde-cross-linking of block Alanine substitutions in the ExbD disordered region. Strains were grown to mid-exponential phase, at which point formaldehyde-cross-linking was performed as described in Materials and Methods. Equivalent numbers of cells were visualized on immunoblots of 13% SDS-polyacrylamide gels with (A) anti-ExbD polyclonal and (B) anti-TonB monoclonal antibodies. cExbD designates chromosomally expressed ExbD from the wild-type strain, W3110. The parent plasmid, pExbD (pKP999) and its corresponding ExbD block Alanine substitutions were expressed in both RA1045 (Δ*exbD*, Δ*tolQR*) [+] and KP1509 (Δ*exbD*, Δ*tolQR,* Δ*tonB*) [-]. (A) Positions of the previously characterized formaldehyde cross-linked PMF-dependent ExbD-TonB complex (red asterisk on immunoblot), the ExbD-ExbB complex, the ExbD-ExbD homodimer and ExbD monomer are shown on the left (5). Mass markers are shown on the right. (B) Positions of the TonB-FepA complex, the TonB-ExbB complex, the PMF-dependent TonB-ExbD complex and TonB monomer are shown. Mass markers are shown on the right. Below each immunoblot are the levels of the corresponding monomer in each sample. The plasmid identities and concentrations of propionate for chromosomal expression are listed in Table S1.

**Figure S2:**
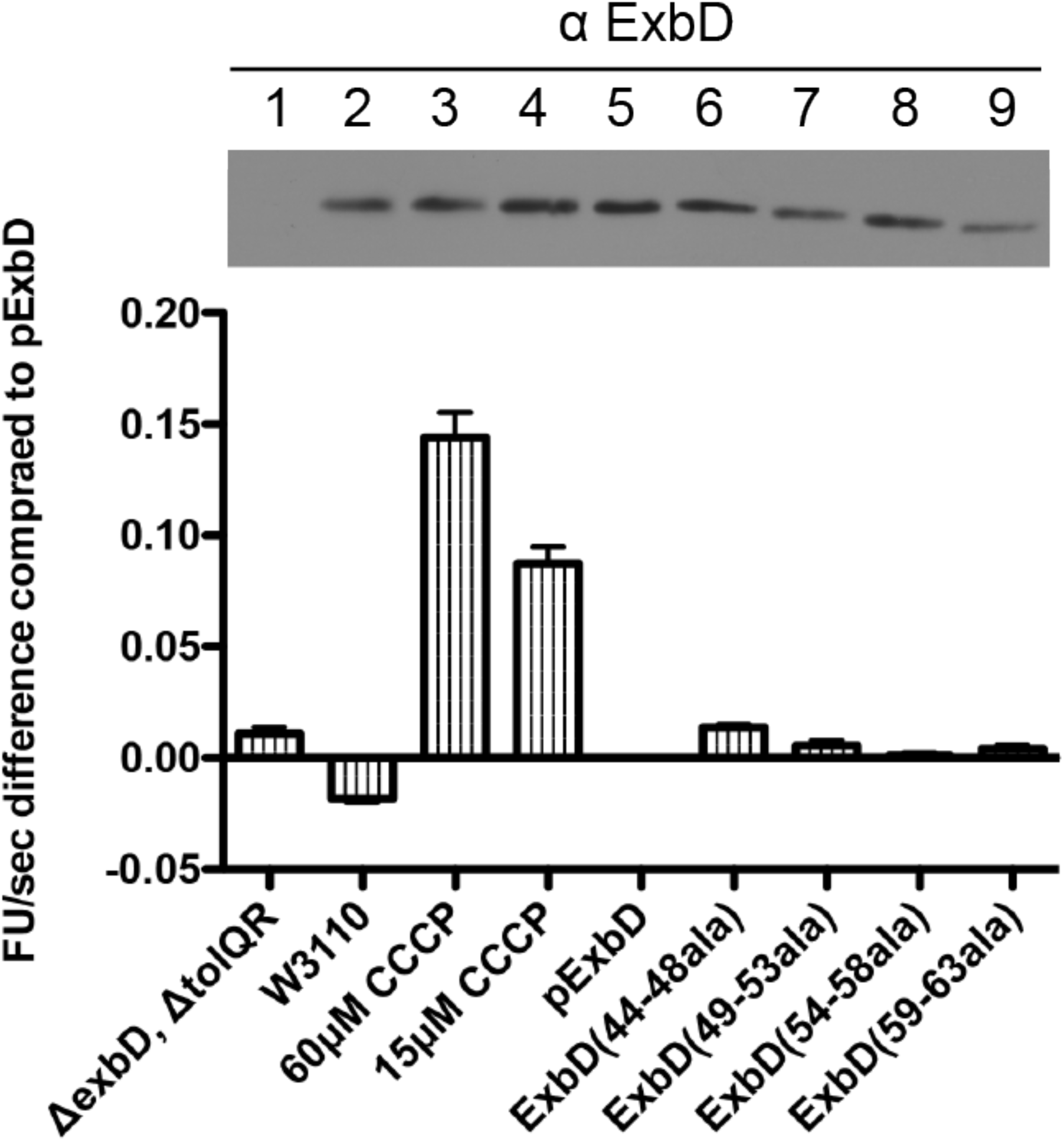
Failure to form the PMF-dependent ExbD-TonB formaldehyde crosslink is not due to loss of PMF; a small, but significant difference is detected among the block Ala substitutions. The rate of ethidium bromide (EtBr) accumulation, an indirect assay for loss of PMF, was measured for ExbD block Alanine substitutions as described in the Materials and Methods. Strains were grown to mid-exponential phase and assayed as described in Methods and Materials. (A) The average EtBr accumulation rates (FU/sec) were calculated relative to the rate of plasmid-encoded ExbD, pExbD (pKP999) expressed at chromosomal levels in RA1045 (Δ*exbD*, Δ*tolQR*). The corresponding steady-state ExbD monomer levels determined by immunoblot analysis from the experiment from one of the trials are shown above the bar graph. cExbD designates chromosomally expressed ExbD from the wild-type strain, W3110. For lanes 3 and 4, W3110 was treated with 60 µM CCCP or 15 µM (as labeled) was prior to addition of EtBr as described in the Materials and Methods. ExbD Alanine substitutions were expressed from plasmids in RA1045 (Δ*exbD*, Δ*tolQR*). The plasmids and concentrations of propionate for chromosomal expression are listed in Table S1. Numbers above the immunoblot correspond to samples below them starting with (Δ*exbD*, Δ*tolQR*) on the left as #1.

**Figure S3:**
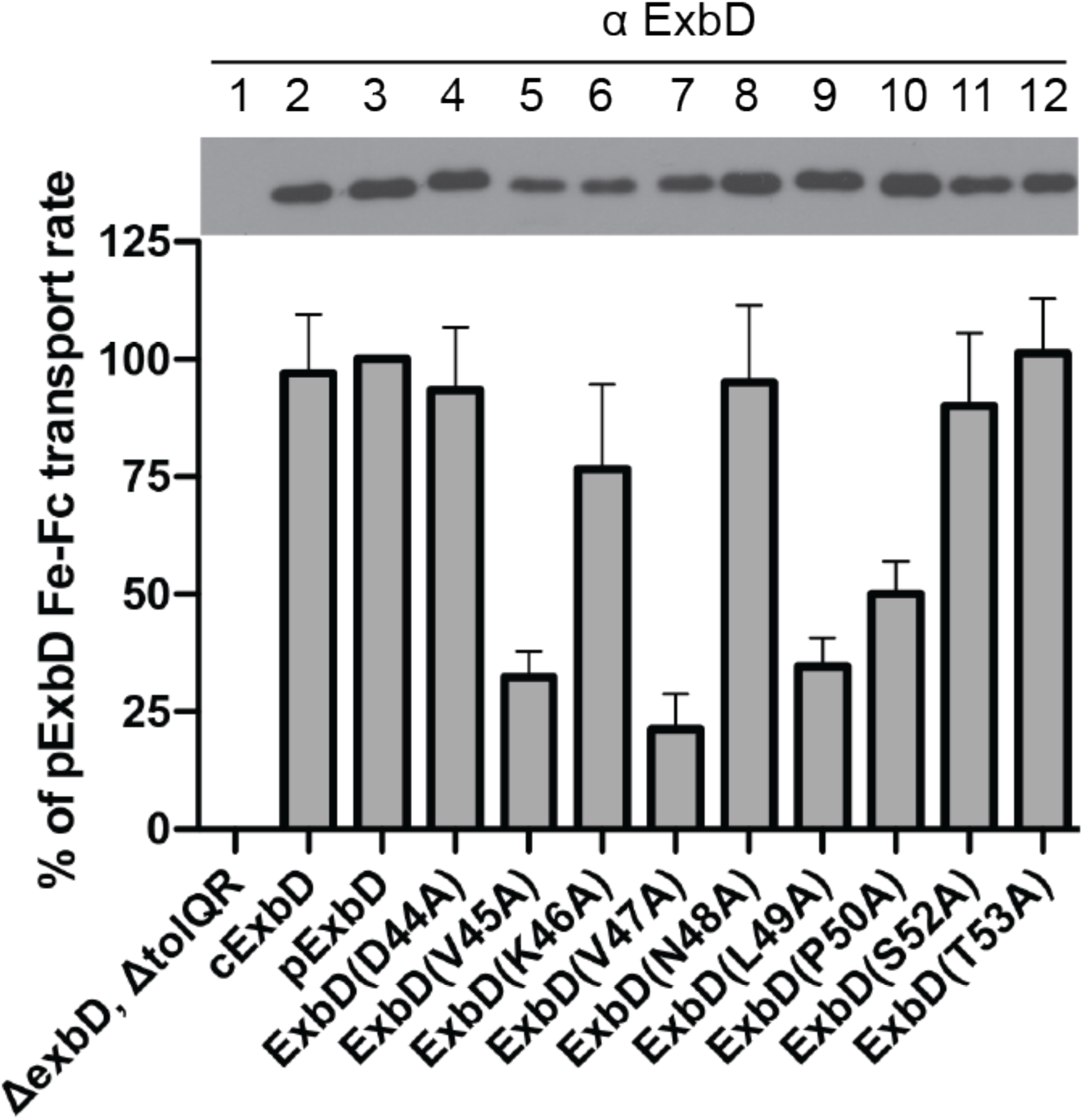
ExbD residues V45, V47, L49 and P50 are important for TonB System activity. Plasmid-encoded wild-type ExbD and Alanine substitution mutants were expressed in RA1045 (Δ*exbD*, Δ*tolQR*). Strains were grown to mid-exponential phase and assayed for the initial rate of ^55^Fe-ferrichrome transport as described in Methods and Materials. The percentages of activity were calculated relative to the initial rate of plasmid-encoded ExbD, pExbD (pKP999). Assays were performed in triplicate and carried out at least twice for each strain. The slopes were determined by a linear regression fit of scintillation counts per minute. The error bars indicate mean ± SEM for a representative triplicate set. cExbD designates chromosomally expressed ExbD from the wild-type strain, W3110. Plasmid identities and concentrations of propionate for chromosomal expression are listed in Table S1. The mean percent of the pExbD ^55^Fe-ferrichrome transport rates for ExbD (V45A), ExbD (V47A), ExbD (L49A), and ExbD (P50A) were 32%, 21%, 34%, and 50% respectively. ExbD residue 51 is naturally an Alanine in *Escherichia coli* W3110 and is reflected in the pExbD initial rate. Steady-state levels of ExbD-specific expression from the transport experiments are indicated above the bar graph.

